# Quantification of vascular networks in photoacoustic mesoscopy

**DOI:** 10.1101/2021.11.22.469541

**Authors:** Emma L. Brown, Thierry L. Lefebvre, Paul W. Sweeney, Bernadette J. Stolz, Janek Gröhl, Lina Hacker, Ziqiang Huang, Dominique-Laurent Couturier, Heather A. Harrington, Helen M. Byrne, Sarah E. Bohndiek

**Affiliations:** Department of Physics, University of Cambridge, JJ Thomson Avenue, Cambridge, CB3 0HE, U.K; Cancer Research UK Cambridge Institute, University of Cambridge, Robinson Way, Cambridge, CB2 0RE, U.K; Mathematical Institute, University of Oxford, Woodstock Road, Oxford, OX2 6GG, U.K

**Keywords:** photoacoustic imaging, vasculature, segmentation, topology

## Abstract

Mesoscopic photoacoustic imaging (PAI) enables non-invasive visualisation of tumour vasculature and has the potential to assess prognosis and therapeutic response. Currently, evaluating vasculature using mesoscopic PAI involves visual or semi-quantitative 2D measurements, which fail to capture 3D vessel network complexity, and lack robust ground truths for assessment of segmentation accuracy. Here, we developed an *in silico*, phantom, *in vivo*, and *ex vivo*-validated end-to-end framework to quantify 3D vascular networks captured using mesoscopic PAI. We applied our framework to evaluate the capacity of rule-based and machine learning-based segmentation methods, with or without vesselness image filtering, to preserve blood volume and network structure by employing topological data analysis. We first assessed segmentation performance against ground truth data of *in silico* synthetic vasculatures and a photoacoustic string phantom. Our results indicate that learning-based segmentation best preserves vessel diameter and blood volume at depth, while rule-based segmentation with vesselness image filtering accurately preserved network structure in superficial vessels. Next, we applied our framework to breast cancer patient-derived xenografts (PDXs), with corresponding *ex vivo* immunohistochemistry. We demonstrated that the above segmentation methods can reliably delineate the vasculature of 2 breast PDX models from mesoscopic PA images. Our results underscore the importance of evaluating the choice of segmentation method when applying mesoscopic PAI as a tool to evaluate vascular networks *in vivo*.

## INTRODUCTION

Tumour blood vessel networks are often chaotic and immature (Brown et al., 2019; Corliss et al., 2019; Hanahan & Weinberg, 2011; Krishna Priya et al., 2016; Nagy & Dvorak, 2012), with inadequate oxygen perfusion and therapeutic delivery (Michiels et al., 2016; Trédan et al., 2007). The association of tumour vascular phenotypes with poor prognosis across many solid cancers (Brown et al., 2019) has generated substantial interest in non-invasive imaging of the structure and function of tumour vasculature, particularly longitudinally during tumour development. Imaging methods that have been tested to visualise the vasculature include whole-body macroscopic methods, such as computed tomography and magnetic resonance imaging, as well as localised methods, such as ultrasound and photoacoustic imaging (PAI) (Brown et al., 2019). Microscopy methods can achieve much higher spatial resolution but are typically depth limited, at up to ∼1mm depth, and frequently applied *ex vivo* (Brown et al., 2019; Jährling et al., 2009; Kelch et al., 2015; Keller & Dodt, 2012; Ntziachristos, 2010).

Of the available tumour vascular imaging methods, PAI is highly scalable and, as such, applicable for studies from microscopic to macroscopic regimes. By measuring ultrasound waves emitted from endogenous molecules, including haemoglobin, following the absorption of light, PAI can reconstruct images of vasculature at depths beyond the optical diffraction limit of ∼1 mm (Beard, 2011; Ntziachristos, 2010; Ntziachristos et al., 2005; Wang & Yao, 2016). State-of-the-art mesoscopic systems now bridge the gap between macroscopy and microscopy, achieving ∼20 μm resolution at up to 3 mm in depth (Omar et al., 2014, 2019). Preclinically, mesoscopic PAI has been used to monitor the development of vasculature in several tumour xenograft models (Haedicke et al., 2020; Omar et al., 2015; Orlova et al., 2019) and can differentiate aggressive from slow-growing vascular phenotypes (Orlova et al., 2019). Studies to-date, however, have been largely restricted to qualitative analyses due to the challenges of accurate 3D vessel segmentation, quantification and robust statistical analyses (Haedicke et al., 2020; Imai et al., 2017; Omar et al., 2015, 2019; Orlova et al., 2019; Rebling et al., 2021). Instead, PAI quantification is typically manual and ad-hoc, with 2D measurements often extracted from 3D PAI data (Haedicke et al., 2020; Imai et al., 2017; Lao et al., 2008; Orlova et al., 2019; Soetikno et al., 2012), reducing repeatability and comparability across datasets.

To assess the performance and accuracy of such vessel analyses, ground truth datasets are needed with *a priori* known features (Krig & Krig, 2014). Creating full-network ground truth reference annotations could be achieved through comprehensive manual labelling of PAI data, but this is difficult due to: the lack of available experts to perform annotation with a new imaging modality; the time taken to label images; and the inherent noise and artefacts present in PAI data. Despite the numerous software packages available to analyse vascular networks (Corliss et al., 2019), their performance in mesoscopic PAI has yet to be evaluated, hence there is an unmet need to improve the quantification of vessel networks in PAI, particularly given the increasing application of PAI in the study of tumour biology (Haedicke et al., 2020; Omar et al., 2019; Orlova et al., 2019).

To quantify PAI vascular images and generate further insights into the role of vessel networks in tumour development and therapy response, accurate segmentation of the vessels must be performed (Corliss et al., 2019) (see step 1 in Figure 1). A plethora of segmentation methods exist and can be broadly split into two categories: rule-based and machine learning-based methods. Rule-based segmentation methods encompass techniques that automatically delineate the vessels from the background based on a custom set of rules (F. Zhao et al., 2019). These methods provide less flexibility and tend to consider only a few features of the image, such as voxel intensity (Haedicke et al., 2020; Orlova et al., 2019; Raumonen & Tarvainen, 2018; Soetikno et al., 2012) but they are easy-to-use, with no training dataset requirements. On the other hand, machine learning-based methods, such as random forest classifiers, delineate vessels based on self-learned features (Moccia et al., 2018; F. Zhao et al., 2019). Nonetheless, learning-based methods are data-driven, requiring large and high-quality annotated datasets for training and can have limited applicability to new datasets. To tackle some of these issues, several software packages have been developed in recent years, and have become increasingly popular in life science research (Berg et al., 2019; Corliss et al., 2019; Sommer et al., 2011). Prior to segmentation, denoising and feature enhancement methods, such as Hessian-matrix based filtering, can also be applied to overcome the negative impact of noise and/or to enhance certain vessel structures within an image (Oruganti et al., 2013; Ul Haq et al., 2016; H. Zhao et al., 2019).

**Figure 1.**
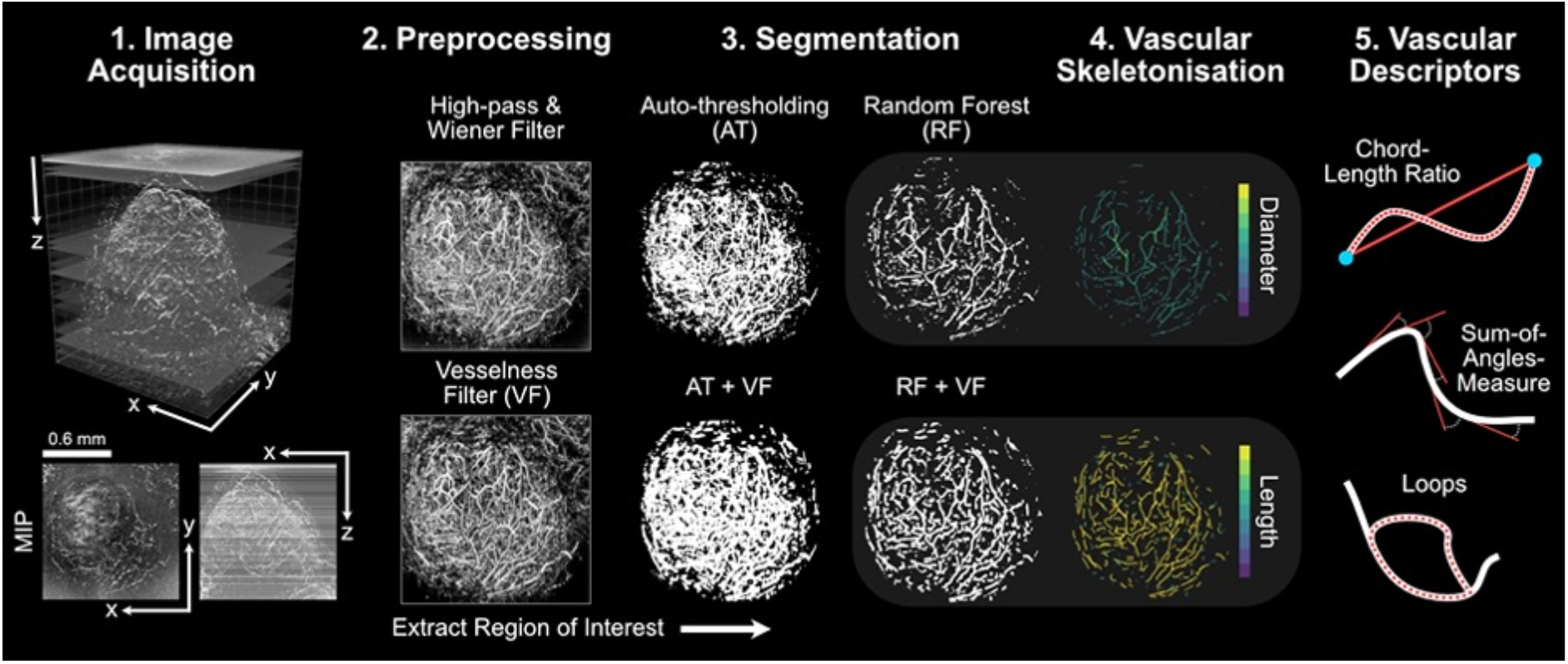
The mesoscopic photoacoustic image analysis pipeline. 1) Images are acquired and reconstructed at a resolution of 20 x 20 x 4 μm^3^ (PDX tumour example shown with axial and lateral maximum intensity projections – MIPs). 2) Image volumes are pre-processed to remove noise and homogenise the background signal (high-pass and Wiener filtering followed by slice-wise background correction). Vesselness image filtering (VF) is an optional and additional feature enhancement method. 3) Regions of interest (ROIs) are extracted and segmentation is performed on standard and VF images using auto-thresholding (AT or AT + VF, respectively) or random forest-based segmentation with ilastik (RF or RF + VF, respectively). 4) Each segmented image volume is skeletonised (skeletons with diameter and length distributions shown for RF and RF + VF, respectively). 5) Statistical and topological analyses are performed on each skeleton to quantity vascular structures for a set of vascular descriptors. All images in steps 2-4 are shown as x-y MIPs.

Here, we establish ground truth PAI data based on simulations conducted using synthetic vascular architectures generated *in silico* and, also using a photoacoustic string phantom, composed of a series of synthetic blood vessels (strings) of known structure, which can be imaged in real-time. Against these ground truths, we compare and validate the performance of two common vessel segmentation methods, with or without the application of 3D Hessian matrix-based vesselness image filtering feature enhancement of blood vessels (steps 2 & 3 in Figure 1). Following skeletonisation of the segmentation masks, we perform statistical and topological analyses to establish how segmentation influences the architectural characteristics of a vascular network acquired using PAI (steps 4 & 5 in Figure 1). Finally, we apply our segmentation and analysis pipeline to two *in vivo* breast cancer models and undertake a biological validation of the segmentation and subsequent statistical and topological descriptors using *ex vivo* immunohistochemistry (IHC). Compared to a rule-based auto-thresholding method, our findings indicate that a learning-based segmentation, via a random forest classifier, is better able to account for the artefacts observed in our 3D mesoscopic PAI datasets, providing a more accurate segmentation of vascular networks. Statistical and topological descriptors of vascular structure are influenced by the chosen segmentation method, highlighting a need to validate and standardise segmentation methods in PAI for increased reproducibility and repeatability of mesoscopic PAI in biomedical applications.

## RESULTS

### *In silico* simulations of synthetic vasculature enable segmentation precision to be evaluated against a known ground truth

Our ground truth consisted of a reference dataset of synthetic vascular network binary masks (n=30) generated from a Lindenmayer System, referred to as L-nets (Figure 2**; Supplementary Movie 1** for 3D visualisation). We simulated PAI mesoscopy data from these L-nets (Figure 2A) and subsequently used vesselness filtering (VF) as an optional and additional feature enhancement method (Figure 2B). The four segmentation pipelines selected for testing (Figure 1) were applied to the simulated PAI data (Figure 2C), that is, all images were segmented with:

1. Auto-thresholding using a moment preserving method (AT);
2. Auto-thresholding using a moment preserving method with vesselness filtering pre-segmentation (AT+VF);
3. Random forest classifier (RF);
4. Random forest classifier with vesselness filtering pre-segmentation (RF+VF).

**Figure 2.**
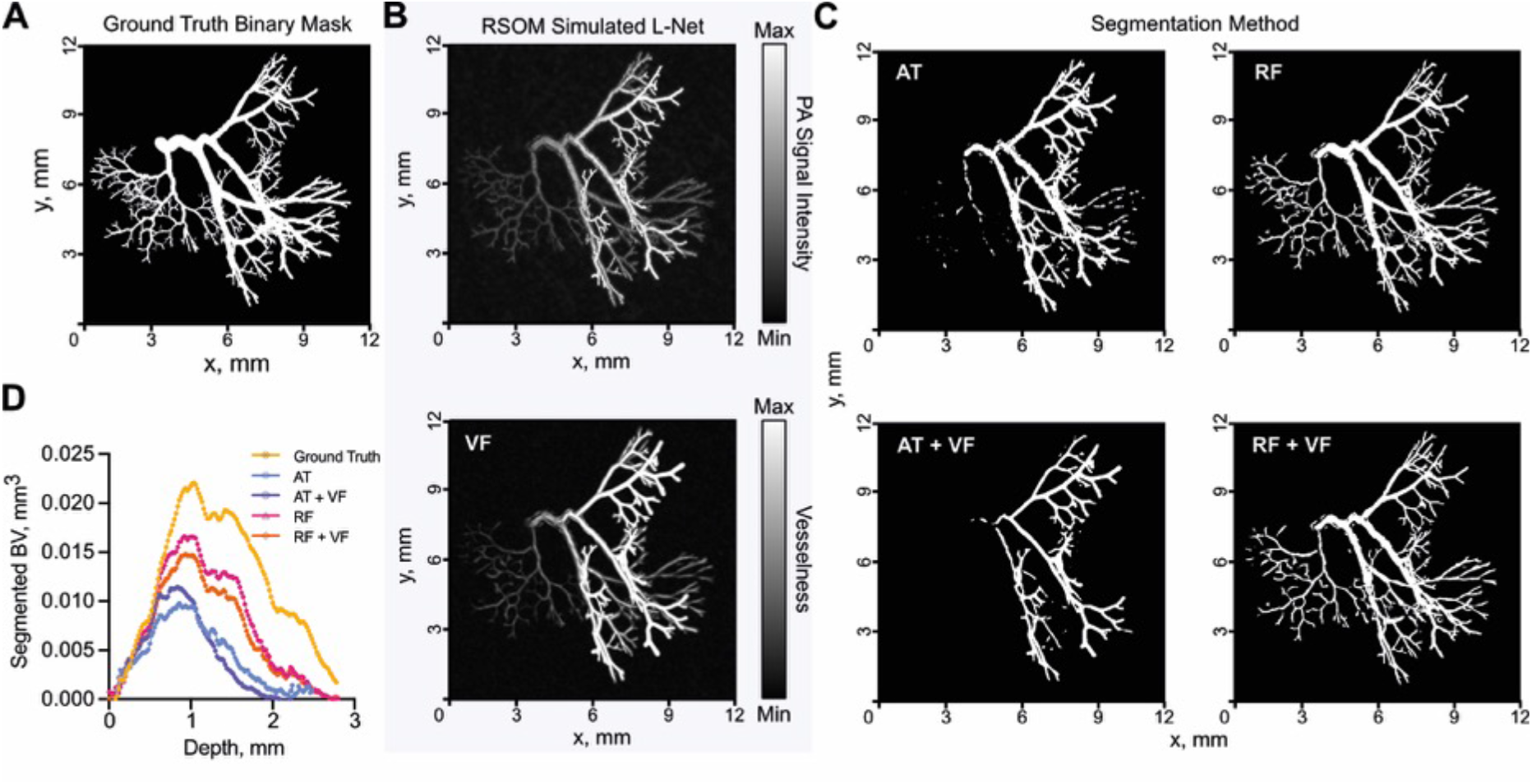
Exemplar vascular architectures generated *in silico* and processed through our photoacoustic image analysis pipeline. (A-C) XY maximum intensity projections of L-net vasculature. (A) Ground truth L-Net binary mask used to simulate raster-scanning optoacoustic mesoscopy (RSOM) image shown in (B, top) and subsequent optional vesselness filterining (VF) (B, bottom). (C) Segmented binary masks generated using either auto-thresholding (AT), auto-thresholding after vesselness filtering (AT + VF), random forest classification (RF); or random forest classification after vesselness filtering (RF+VF). (D) Segmented blood volume (BV) average across L-net image volumes, plotted against image volume depth (mm). For (D) n=30 L-nets. See Supplementary Movie 1 for 3D visualisation.

Visually, RF methods appear to segment a larger portion of synthetic blood vessels (Figure 2C) and they are particularly good at segmenting vessels at depths furthest from the simulated light source (Figure 2D). A key image quality metric in the context of segmentation is the signal-to-noise (SNR), which is degraded at greater depth (Figure 3A). To evaluate the relative performance of the methods, we compared the segmented and skeletonised blood volumes (BV) from the simulated PAI data to the known ground truth from the L-net. Here, we found that the learning-based RF segmentation outperformed the others in making the segmentation masks, with significantly higher R^2^ (segmented BV: AT: 0.68, AT+VF: 0.58, RF: 0.84, RF+VF: 0.89, Figure 3B skeleton BV: AT: 0.59, AT+VF: 0.73, RF: 0.90, RF+VF: 0.93, Figure 3C) and lower mean-squared error (MSE) (Figure 3D), with respect to the ground truth L-net volumes, compared to both AT methods (p<0.0001 for all comparisons). Bland-Altman plots, which we used to illustrate the level of agreement between segmented and ground truth vascular volumes, showed a mean difference compared to the reference volume of 0.61 mm^3^ (limits of agreement, LOA -0.48 to 1.7 mm^3^, Figure 3E) and F1 score of 0.73 ± 0.11 (0.49-0.88) for RF segmentation, albeit with a wide variation indicated by the LOA. RT+VF segmentation resulted in a similar mean difference 0.74 mm^3^ (LOA -0.50 to 2.0 mm^3^, Figure 3F) and F1 score of 0.66 ± 0.11 (0.44-0.84). In comparison, the rule-based AT segmentation showed poor performance in segmenting vessels at depth (Figure 2C**, Supplementary Movie 1**), yielding a mean difference of 1.1 mm^3^ (LOA -0.60 to 2.8 mm^3^) and as with RT+VF, AT+VF did not improve the result, yielding the same mean difference of 1.1 mm^3^ (LOA -0.52 to 2.8 mm^3^) (Figure 3G,H). F1 scores were poor for both AT methods, with 0.39 ± 0.10 (0.21-0.59) for AT and 0.37 ± 0.09 (0.16-0.52) for AT+VF.

**Figure 3.**
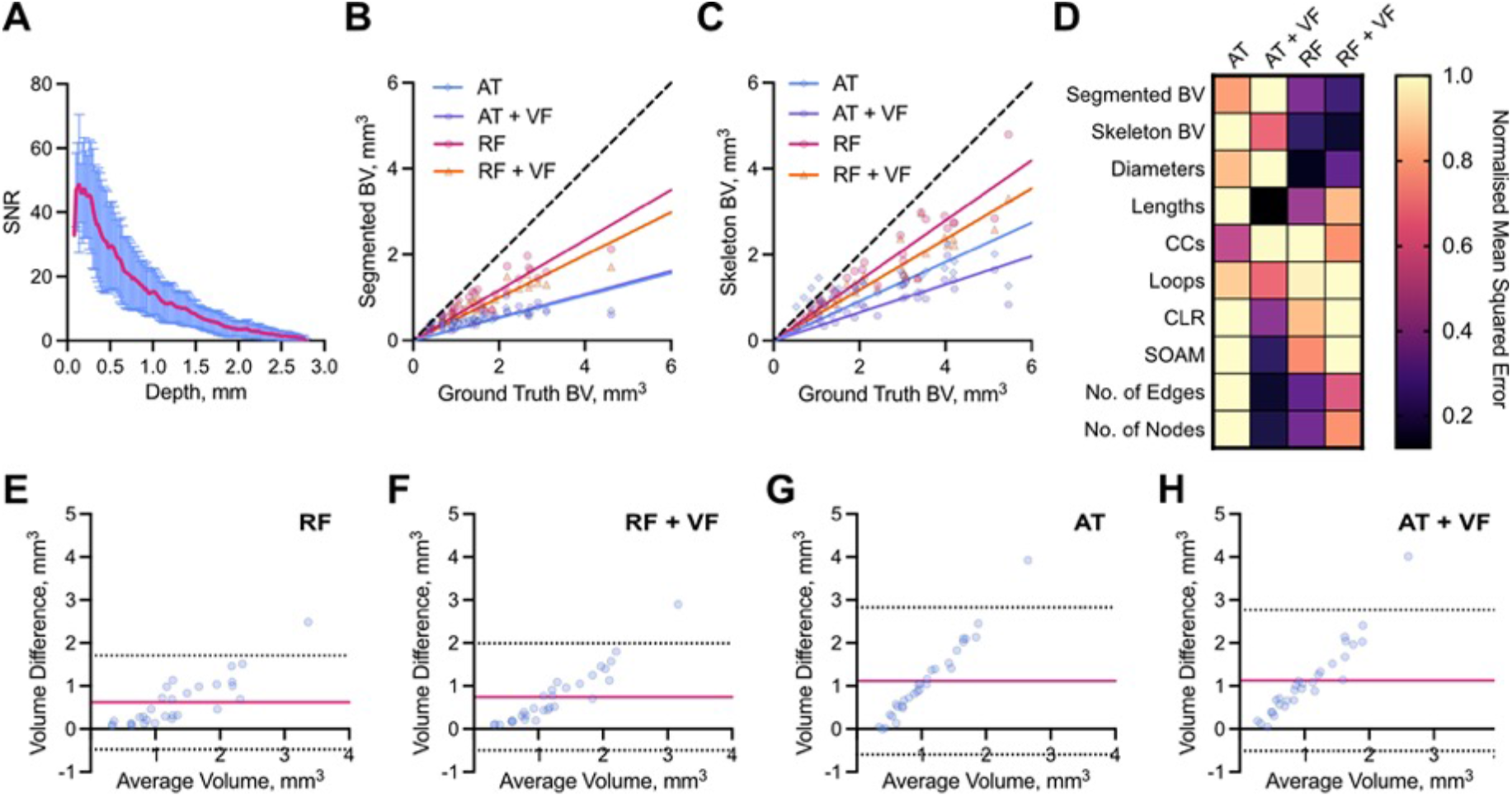
Learning-based random forest classifier outperforms rule-based auto-thresholding in segmenting simulated PAI vascular networks. (A) Depth-wise comparison of signal-to-noise ratio (SNR) measured in PAI-simulated L-nets across depth. (B,C) A comparison between ground truth blood volume (BV) and (B) segmented or (C) skeletonised blood volumes (BV). The dashed line indicates a 1:1 relationship. (D) Heat map displaying normalised (with respect to the maximum of each individual descriptor) mean-squared error comparing our vascular descriptors, calculated from segmented and skeletonised L-nets compared to ground truth L-nets, to each segmentation method. Abbreviations defined: connected components, β0 (CC), chord-to-length ratio (CLR), sum-of-angle measure (SOAM). (E-H) Bland-Altman plots comparing blood volume measurements from ground truth L-nets with that of each segmentation method: (E) RF, (F) RF+VF, (G) AT, (H) AT+VF. Pink lines indicate mean difference to ground truth, whilst dotted black lines indicate limits of agreement (LOA). For all subfigures n=30 L-nets.

In all cases, the mean difference shown in Bland-Altman plots increased with ground truth vascular volume, especially in the rule-based AT segmentation, which would be expected due to the restricted illumination geometry of photoacoustic mesoscopy. Since more vessel structures lie at a greater distance from the simulated light source in larger L-nets, they suffer from the depth-dependent decrease in SNR (Figure 3A). RF segmentation was better able to cope with the SNR degradation, particularly at distances beyond ∼1.5 mm, compared to the AT segmentation, which consistently underestimated the vascular volume.

Next, we skeletonised each segmentation mask to enable us to perform statistical and topological data analysis (TDA) to test how each segmentation method quantitatively influences a core set of vessel network descriptors (Stolz et al., 2020). These descriptors allowed us to evaluate the performance of the different segmentation methods in respect of the biological characterisation of the tumour networks. We used the following statistical descriptors: vessel diameters and lengths, vessel tortuosity (sum-of-angles measure, SOAM) and vessel curvature (chord-to-length ratio, CLR). Our topological network descriptors are connected components (Betti number β_0_) and looping structures (1D holes, Betti number β_1_) (see Supplementary Table 1 for descriptor descriptions).

Here, the accuracy and strength of relationship between the segmented and ground truth vascular descriptors, calculated by MSE (see Figure 3D) and R^2^ values (Supplementary Figure 1A-I) respectively, gave the same conclusions. Across all skeletons, we measured an increased number of connected components (β_0_) and changes to the number of looping structures (β_1_) from the simulated compared to the ground truth L-nets, resulting in low R^2^ and high MSE for all methods (Figure 3D). The observed changes in these topological descriptors arise due to depth-dependent SNR and PAI echo artefacts. For all other descriptors, AT+VF outperformed the other segmentation methods in its ability to accurately preserve the architecture of the L-nets, with higher R^2^ and lowest MSE values for vessel lengths, CLR, SOAM, number of edges and number of nodes (Figure 3D).

Vessel diameters are accurately preserved by both RF segmentation methods, supporting our observation that these methods perform accurate vascular volume segmentation. We note that the number of edges and nodes are also well preserved by RF and RF+VF. This further supports the high accuracy of both RF methods to segment vascular structures.

### Random forest classifier accurately segments a string phantom

We next designed a phantom test object (Supplementary Figure 2) to further compare the performance of our segmentation pipelines in a ground truth scenario. Agar phantom images (n=7) were acquired using a photoacoustic mesoscopy system and contained three strings of the same known diameter (126 μm), length (∼8.4 mm) and consequently volume (104.74 μm^3^), positioned at 3 different depths, 0.5 mm, 1 mm, and 2 mm, respectively (Figure 4A,B**; Supplementary Movie 2**). Consistent with our *in silico* experiments, the accuracy of skeletonised string volumes decreased as a function of depth across all methods (Figure 4C), due to the decreased SNR with depth (Figure 4D). Interestingly, the significance of this decrease was very high for all comparisons (top vs. middle, top vs. bottom and middle vs. bottom) in both AT methods (all p<0.001), but we observed an improvement in string volume predictions across depth for both RF methods, such that middle vs. bottom string volumes were not significantly different in RF+VF (p=0.42).

**Figure 4.**
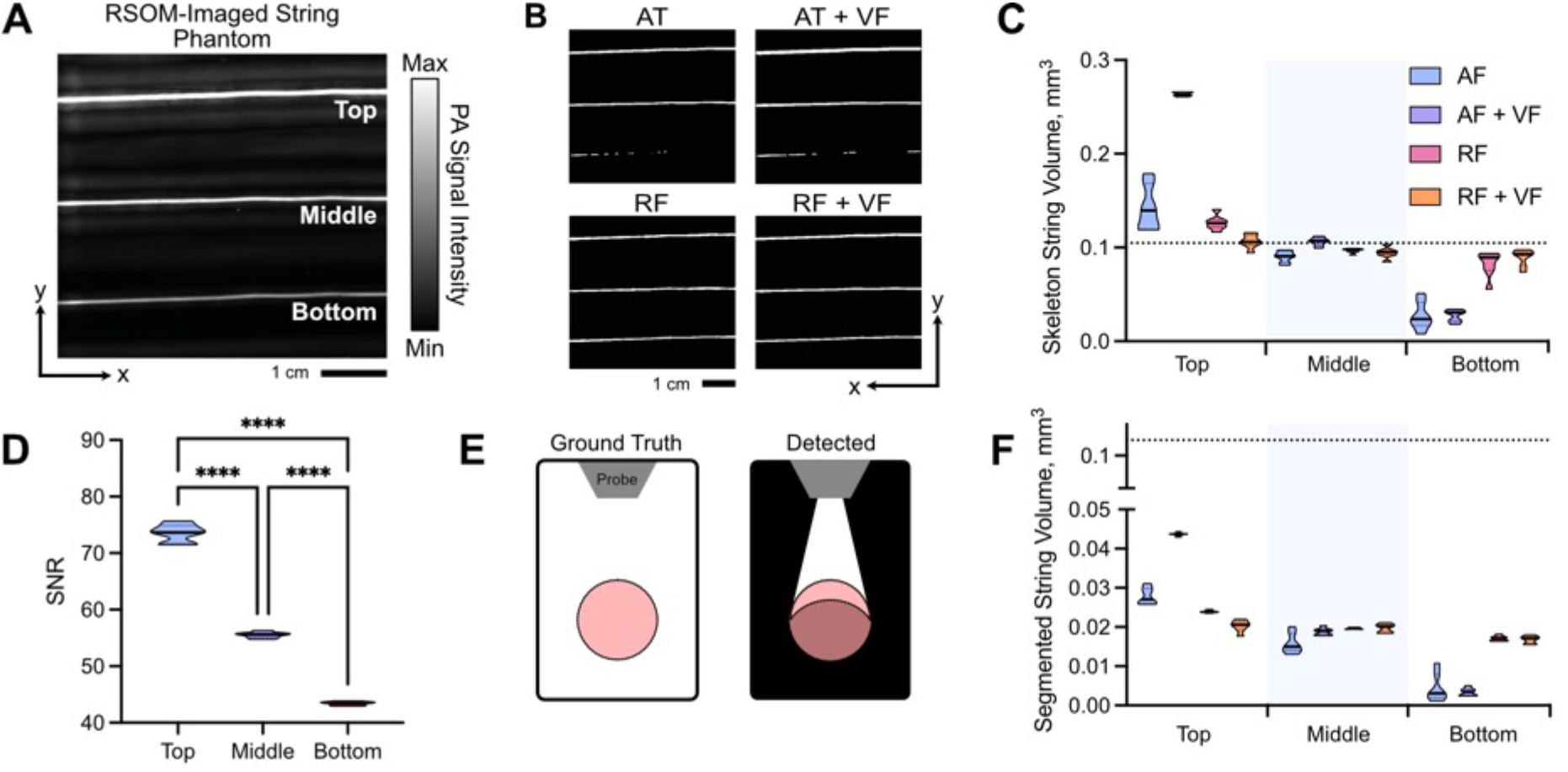
Random forest classifier outperforms auto-thresholding in segmenting a string phantom. XY maximum intensity projections of string phantom imaged with RSOM show that random forest-based segmentation outmatches auto-thresholding when correcting for depth-dependent SNR. (A) Photoacoustic mesoscopy (RSOM) image shows measured string PA signal intensity with top (0.5 mm), middle (1 mm) and bottom (2 mm) strings labelled. (B) Binary masks are shown following segmentation using: (AT) auto-thresholding; (RF) Random forest classifier; (AT+VF) vesselness filtered strings with auto-thresholding; and (RF+VF) vesselness filtered strings with random-forest classifier. (C) Skeletonised string volume calculated from segmented images of 3 strings placed at increasing depths in an agar phantom. Results from all 4 segmentation pipelines are shown. All volume comparisons (top vs. middle, top vs. bottom, middle vs. bottom) where significant (p<0.05) except middle vs. bottom for RF+VF (p=0.42). (D) SNR decreases with increasing depth. (E) Illumination geometry: known cross-section of string outlined (left); during measurement, signal is detected from the partially illuminated section (outlined) resulting in an underestimation in string volume (right). (F) String volume calculated pixel-wise from the segmented binary mask. (C,D,F) Data represented by truncated violin plots with interquartile range (bold) and median (dotted), ****=p<0.0001 (n=7 scans). (C,F) Dotted line indicates ground truth volume 0.105 mm^3^. See Supplementary Movie 2 for 3D visualisation.

The illumination geometry of the photoacoustic mesoscopy system means that vessels or strings are underrepresented when detected as the illumination source is located at the top surface of the tissue or phantom (Figure 4E). As a result, all string volumes computed from the segmented images are inaccurate relative to ground truth suggesting that blood volume cannot be accurately predicted from segmented PA images (Figure 4F). Skeletonisation provides a more accurate prediction of vessel and string volume as it approximates the undetected section by representing these objects as axisymmetric tubes (Figure 4C,F).

### Vesselness filtering of in vivo tumour images impacts computed blood volume

Having established the performance of our AT- and RF-based segmentation methods *in silico* and in a string phantom, next we sought to determine the influence of the chosen method in quantifying tumour vascular networks from size-matched breast cancer patient-derived xenograft (PDX) tumours of two subtypes (ER-*n*=6; ER+ *n*=8, total *n*=14).

Visual inspection of the tumour networks subjected to our processing pipelines suggests that VF increases vessel diameters *in vivo* (Figure 5A-C**; see Supplementary Movie 3** for 3D visualisation). This could be due to acoustic reverberations observed surrounding vessels *in vivo*, which VF scores with high vesselness, spreading the apparent extent of a given vessel and ultimately increased volume. Our quantitative analysis confirmed this hypothesis, where significantly higher skeletonised blood volumes were calculated in the AT+VF and RF+VF masks compared to AT and RF alone (Figure 5D).

**Figure 5.**
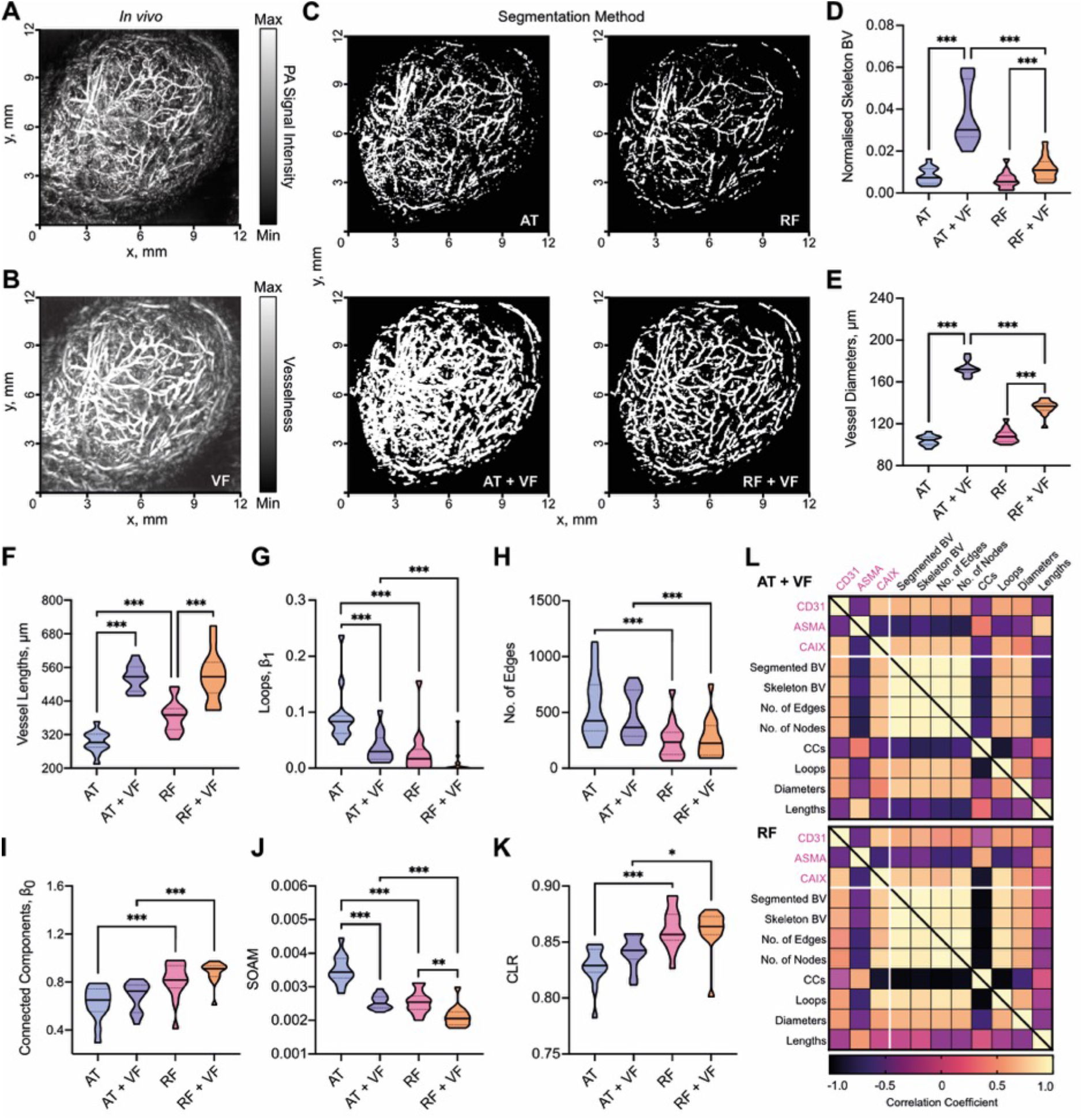
Vesselness filtering increases blood volume calculations from *in vivo* tumour images. XY Maximum intensity projections of breast PDX tumours imaged with RSOM: (A) original image before segmentation; (B) original image with vesselness filtering (VF) applied; (C) a panel showing segmentation with each method (AT: auto-thresholding, AT+VF: auto-thresholding with VF, RF: random forest classifier, and RF + VF: random forest with VF). (D) Skeletonised tumour blood volume (BV) from all 4 segmentation methods normalised to ROI volume. Statistical and topological data analyses were performed on skeletonised tumour vessel vascular networks for the following descriptors: (E) Total number of edges; (F) Connected components normalised by network volume, β0; (G) loops normalised by network volume, β1; (H) sum-of-angle measure (SOAM); (I) vessel lengths; (J) vessel diameters; (K) chord-to-length ratio (CLR). In (D-K), data are represented by truncated violin plots with interquartile range (dotted) and median (bold). Pairwise comparisons of AT vs. AT+VF, AT vs. RF, RF vs. RF+VF and AT+VF vs. RF+VF calculated using a linear mixed effects model (*= p<0.05, **=p<0.01, ***=p<0.001,). L) Matrix of correlation coefficients for comparisons between IHC, BV and vascular descriptors for (top) AT+VF and (bottom) RF segmented networks. Pearson or spearman coefficients are used as appropriate, depending on data distribution. For (D) n=14, (E-K) n=13 due to imaging artefact in one image which will impact our vascular descriptors. For (L) comparisons involving BV n=14, all other vascular descriptors n=13. See Supplementary Movie 3 for 3D visualisation.

### Network structure analyses and comparisons to *ex vivo* immunohistochemistry of tumour vasculature are impacted by the choice of segmentation method

Next, we computed vascular descriptors for our dataset of segmented *in vivo* images. As expected from our initial *in silico* and phantom evaluations, VF led to increased vessel diameters and lengths (Figure 5E,F), as well as blood volume. Our *in silico* analysis indicated that AT performs poorly in differentiating vessels from noise and introduces many vessel discontinuities (Supplementary Table 1). This was exacerbated *in vivo* where more complex vascular networks and real noise lead to an increase in segmented blood volume (p<0.01), looping structures (Figure 5G), a greater number of edges (Figure 5H), and reduced number of connected components (Figure 5I).

Our prior *in silico* and phantom experiments indicate that RF-based methods have a greater capacity to segment vessels at depth. Similarly, we observe more connected components for RF-based methods *in vivo* (Figure 5I) along with lower SOAM (Figure 5J) and higher CLR (Figure 5K), suggesting that RF-segmented vessels have reduced tortuosity and curvature compared to AT+VF segmented vessels. These *in vivo* findings support our observations from *in silico* and phantom studies where RF-based methods provide the most reliable prediction of vascular volume, whereas AT+VF best preserves architecture towards the tissue surface.

Next, we sought to assess how our vascular metrics correlated with the following *ex vivo* IHC descriptors: CD31 staining area (to mark vessels), ASMA vessel coverage (as a marker of pericyte/smooth muscle coverage and vessel maturity) and CAIX (as a marker of hypoxia) to provide *ex vivo* biological validation of our *in vivo* descriptors. Our *in silico*, phantom and *in vivo* analyses indicate that AT+VF and RF are the top performing segmentation methods and so we focussed on these (results for AT and RF+VF can be found in Supplementary Figure 3). We note that none of the vascular metrics derived from AT segmented networks correlated with IHC descriptors.

Both AT+VF and RF skeletonised blood volume correlate with CD31 staining area (r=0.54, p=0.05; and r=0.61, p=0.02 respectively; Figure 5L). This is as expected as elevated CD31 indicates a higher number of blood vessels and, consequently, higher vascular volume. The following correlations are observed for ASMA vessel coverage: vessel diameters (r=-0.41, p=0.17; and r=-0.43, p=0.14, respectively); looping structures (r=-0.68, p=0.01; and r=-0.58, p=0.04, respectively); number of edges (r=-0.69, p=0.01; and r=-0.65, p=0.02, respectively); number of nodes (r=-0.70, p=0.01; and r=-0.65, p=0.02, respectively); vessel lengths (r=0.76, p=0.03; and r=0.5, p=0.08, respectively); connected components (r=0.38, p=0.22; and r=0.59, p=0.03, respectively). Considering the strengths of AT+VF and RF, these results are biologically intuitive as tumour vessel maturation may lead to higher pericyte coverage, lower vessel density and the pruning of redundant vessels. Elevated pericyte coverage is known to decrease vessel diameters (Barlow et al., 2013), whereas high vessel density resulting from high angiogenesis rates can result in immature vessel networks (Brown et al., 2019). Pruning may lead to a reduction in looping structures and, consequently, an increase in vessel lengths or vascular subnetworks.

Finally, levels of hypoxia in the tumours, measured by CAIX IHC, positively correlated in both AT+VF and RF methods with skeletonised blood volume (r=0.72, p=0.007; and r=0.72, p=0.004, respectively), number of edges (r=0.59, p=0.04; and r=0.84, p<0.001, respectively), nodes (r=0.72, p=0.007; and r=0.84, p<0.001, respectively) and looping structures (r=0.61, p=0.03; and r=0.85, p<0.001, respectively). In the case of blood volume, edges and nodes, these results are expected as it has been shown that breast cancer tumours with dense but immature and dysfunctional vasculatures exhibit elevated hypoxia(Brown et al., 2019; Quiros-Gonzalez et al., 2018), likely due to poor perfusion. CAIX negatively correlated with connected components for RF networks (r=-0.87, p<0.001) (Figure 5L), reflecting results for ASMA vessel coverage. Our cross-validation between *ex vivo* IHC and vascular descriptors indicate that RF and AT+VF segmentation methods can reliably capture biological characteristics in tumours.

### Ex vivo immunohistochemistry and network structural analyses highlight distinct vascular networks between ER- and ER+ breast patient-derived xenograft tumours

Finally, we quantified and compared IHC and our vascular descriptors between the two breast cancer subtypes represented (RF in Figure 6; AT+VF in Supplementary Figure 4; similar trends and significances are observed unless stated otherwise). From analysis of IHC images (Figure 6A), ER-tumours had higher CD31 staining area (Figure 6B), poorer ASMA+ pericyte vessel coverage (Figure 6C) and higher CAIX levels (Figure 6D) compared to ER+ tumours. Our IHC data supports our RF-derived vascular descriptors, where we found that ER-tumours had denser networks, with higher blood volume, diameter and looping structures (Figure 6E,F**,G**). ER+ tumours have a sparse network but showed more subnetworks (Figure 6H) with significantly longer vessels in AT+VF segmented networks (p<0.05, Supplementary Figure 4C), which could indicate a more mature vessel network based on our prior correlative analyses. No significant differences between the two models were observed for blood vessel tortuosity and curvature (Figure 6J,K).

**Figure 6.**
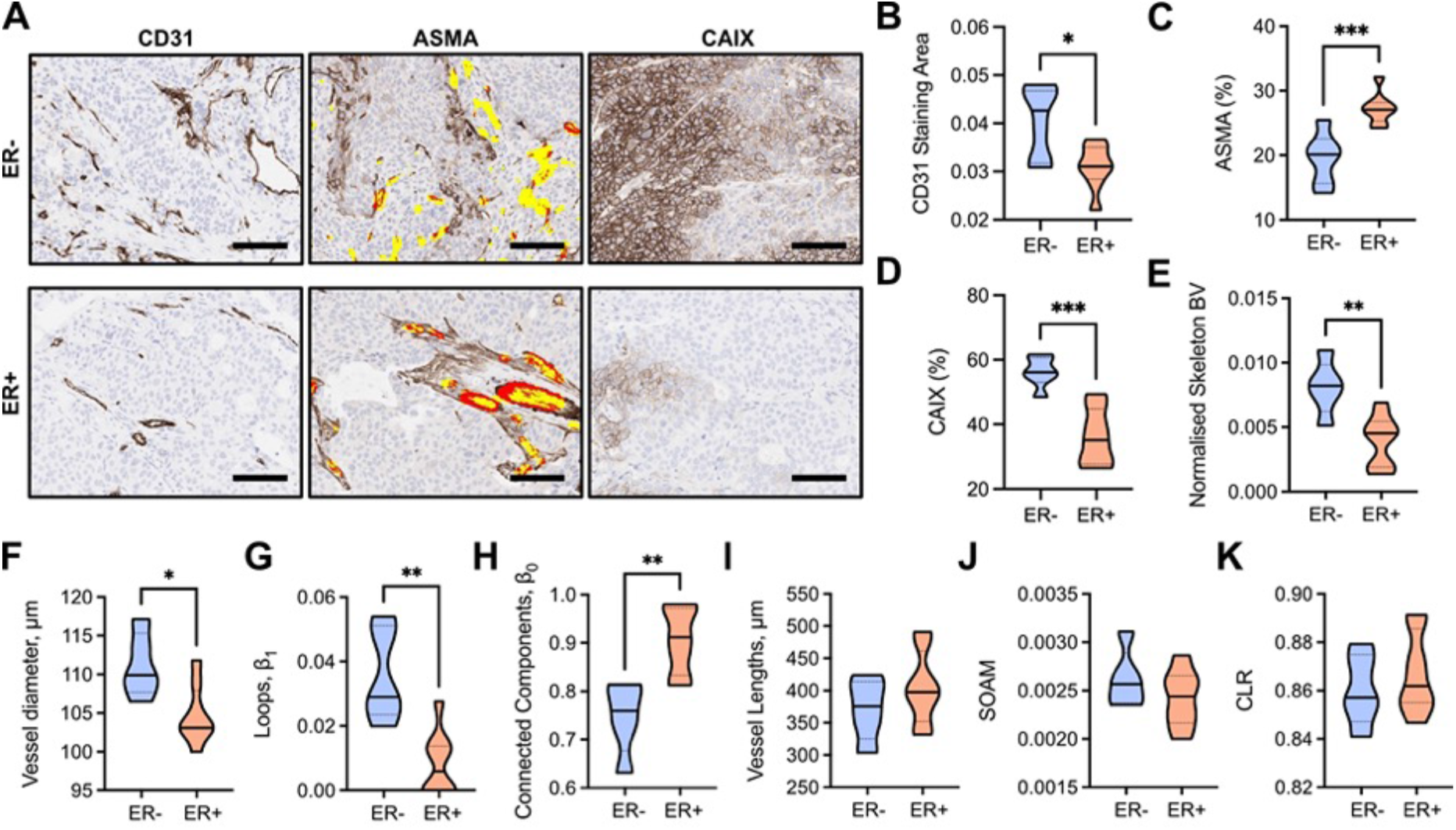
ER-PDX tumours have dense and immature vascular networks which result in hypoxic tumour tissue. (A) Exemplar IHC images of CD31, ASMA and CAIX stained ER- and ER+ tumours. Scale bar=100µm. Brown staining indicates positive expression of marker. ASMA sections display CD31 overlay, where red indicates areas where CD31 and ASMA are colocalised (ASMA vessel coverage) and yellow indicates areas where CD31 is alone. (B) CD31 staining area quantified from CD31 IHC sections and normalised to tumour area. (C) ASMA vessel coverage of CD31+ vessels (number of red pixels/number of red+yellow pixels, expressed as a percentage) on ASMA IHC sections. (D) CAIX total positive pixels as a percentage of the total tumour area pixels on CAIX IHC sections. (E-K) Statistical and topological data analyses comparing ER- and ER+ tumours. Data are represented by truncated violin plots with interquartile range (dotted black) and median (solid black). Comparisons between ER- and ER+ tumours made with unpaired t-test. *= p<0.05, **=p<0.01, ***=p<0.001. For (B-E) ER-n=6, ER+ n=8. For (F-K) ER-n=5, ER+ n=8, one ER-image excluded with artefact that would impact the measured vascular descriptors.

## DISCUSSION

Mesoscopic PAI enables longitudinal visualisation of blood vessel networks at high resolution, non-invasively and at depths beyond the optical diffraction limit of 1 mm (Ntziachristos, 2010; Ntziachristos et al., 2005; Omar et al., 2019; Wang & Yao, 2016). To quantify the vasculature, PA images need to be accurately segmented. Manual annotation of vasculature in 3D PAI is difficult due to depth-dependent signal-to-noise and imaging artefacts. Whilst a plethora of vascular segmentation techniques are available (Corliss et al., 2019; Moccia et al., 2018), their application in PAI has been limited due to a lack of an available ground truth for comparison and validation.

In this study, we first sought to address the need for ground truth data in PAI segmentation. We generated two ground truth datasets to assess the performance of rule-based and machine learning-based segmentation approaches with or without feature enhancement via vesselness filtering. The first is an *in silico* dataset where PAI was simulated on 3D synthetic vascular architectures; the second is an experimental dataset acquired from a vessel-like string phantom. These allowed us to evaluate the ability of different segmentation methods to preserve blood volume and vascular network structure.

Our first key finding is that machine learning-based segmentation using RF classification provided the most accurate segmentation of vessel volumes across our *in silico*, phantom and *in vivo* datasets, particularly at depths beyond ∼1.5mm, where SNR diminishes due to optical attenuation. Compared to the AT approaches, RF-based segmentation partially overcomes the depth dependence of PAI SNR since it identifies and learns edge and texture features of vessels at different scales and contrasts. Such intrinsic depth-dependent limitations are often ignored in the literature, where analyses are typically performed on 2D maximum intensity projections for simplicity (Haedicke et al., 2020; Imai et al., 2017; Lao et al., 2008; Omar et al., 2015; Orlova et al., 2019; Soetikno et al., 2012), suggesting that a fully 3D machine learning-based segmentation is needed to accurately recapitulate the complexity of *in vivo* vasculatures measured using PAI.

As blood vessel networks can be represented as complex, interconnected graphs, we performed statistical and topological data analyses (Chung et al., 2019; Stolz et al., 2020) to further assess the strengths and weaknesses of our chosen segmentation methods.

Our second key finding is that AT methods struggle to segment vessels with low SNR, but adding VF outperforms all other methods in preserving vessel lengths, loops, curvature and tortuosity. Additionally, where intensity varies across a vessel structure, this results in many disconnected vessels when segmenting with AT alone, as only the highest intensity voxels will pass the threshold. Only when vesselness filtering is applied does AT do well at preserving topology. VF alters the intensity values from a measure of PA signal to a prediction of ‘vesselness’, generating a more homogeneous intensity across the vessel structures and ultimately a more continuous vessel structure to segment. This likely explains why AT+VF best preserves vessel length and, subsequently, network structure, while AT alone performs poorly. For AT, VF improved BV predictions *in silico* via better preservation of lengths but not diameters, as our phantom experiments indicate that AT+VF overestimates diameter.

Owing to the homogenous intensity of vessels introduced by VF, one could therefore assume that RF+VF would be the most accurate method at preserving network structure (by combining the machine-learning accuracy of segmentation with the shape enhancement of VF). However, this is not the case: RF alone can account for discontinuities in vessel intensity, unlike AT, meaning it does not rely on VF to enhance structural preservation, which is our third key finding. In fact, the slight inaccuracy in diameter preservation introduced by VF *in silico* appears to decrease topology preservation in RF+VF compared to RF alone. As expected, all methods led to an increase in the number of subnetworks (connected components) *in silico*, as these segmentation methods cannot reconnect vessel subnetworks that were disconnected due to poor SNR or imaging artefacts. Given the better segmentation at depth by RF-methods, we hypothesise that these increasingly small subnetworks might have biased the segmentations to underperform in our vascular descriptors. This could be explored in future work, for example, by developing string phantoms with more complex topologies.

Taken together, our results suggest that RF performs feature detection across scales in the manually labelled voxels to learn discriminating characteristics for vessel classification and segmentation. Adding VF before RF segmentation may confound this segmentation framework, because VF systematically smooths images and removes non-cylindrical raw image information, which may have been vital in the RF learning of vascular structures on the training dataset.

Applying statistical and topological analyses to our *in vivo* tumour PDX dataset we observed trends consistent with our *in silico and* phantom experiments. Cross-validating our vascular descriptors with *ex vivo* IHC confirmed that we can extract biologically relevant information from mesoscopic PA images. For example, predictions of BV correlated with endothelial cell and hypoxia markers via CD31 and CAIX staining, respectively; and descriptors relating to the maturation of vascular structures correlated with ASMA vessel coverage. Applying our segmentation pipeline to compare ER- and ER+ breast cancer PDX models showed that descriptors of network structure can capture the higher density and immaturity of ER-vessel networks which result in decreased oxygen delivery and high hypoxia levels in comparison to ER+ tumours.

While our pipeline yields encouraging correlations to the underlying tumour vasculature, avenues of further development exist to: improve the realism of our ground truth data, including advances in simulation complexity, and tissue-specific synthetic and phantom vasculatures. While our *in silico* PAI dataset incorporated the effects of depth-dependent SNR and gaussian noise found in *in vivo* PAI mesoscopic data, further development of the optical simulations could, for example, recapitulate the raster-scanning motion of illumination optical fibres, instead of approximating a simultaneous illumination plane of single-point sources. The limited aperture of the raster-scanning ultrasound transducer could not be simulated in k-Wave as it is not yet implemented for 3D structures. In terms of vascular complexity, our string phantom represents a highly idealised vessel networks but future work could introduce more complex and interconnected vessel-like networks in order to replicate more realistic vascular topologies (Dantuma et al., 2019). Our *ex vivo* IHC descriptors were used to confirm our *in vivo* tumour analyses but did not exhibit correlations across all vascular descriptors. This may be expected as the 2D IHC analysis does not fully encompass the 3D topological characteristics of the vascular network. 3D IHC, microCT or light sheet fluorescence microscopy may provide improved *ex vivo* validation using exogenous labelling to identify 3D vascular structures, such as tortuosity, at endpoint (Epah et al., 2018; Hlushchuk et al., 2019). It should also be noted that we cannot discount the effect of unconscious biases on segmentation performance when manually labelling images with and without VF to train the classifier. The segmentation accuracy of classifiers trained by multiple users could be explored in future work to formally investigate these effects.

Furthermore, the past decade has seen the rise of a multitude of blood vessel segmentation methods using convolutional neural networks and deep learning (Jia & Zhuang, 2021). Applying deep learning to mesoscopic PAI could provide a means to overcome several equipment-related limitations such as: vessel discontinuities induced by breathing motion *in vivo*; vessel orientation relative to the ultrasound transducer; shadow and reflection artefacts; or underestimation of vessel diameter in the z-direction due to surface illumination. Whilst we found that skeletonisation addressed diameter underestimation and observed the influence of discontinuities on the extracted statistical and topological descriptors, they were not deeply characterised or corrected. Nonetheless, whilst deep learning may provide superior performance when fine-tuned to specific tasks, the resulting methods may lack generalisability across tissues with differing SNR and blood structures, requiring large datasets for training. In this study we chose to use software that is open-source and widely accessible to biologists in the life sciences. We believe that such a platform shows more potential to be employed widely with limited computational expertise.

In summary, we developed an *in silico*, phantom*, in vivo*, and *ex vivo-*validated end-to-end framework for the segmentation and quantification of vascular networks captured using mesoscopic PAI. We created *in silico* and string phantom ground truth PAI datasets to validate segmentation of 3D mesoscopic PA images. We then applied a range of segmentation methods to these and images of breast PDX tumours obtained *in vivo*, including cross-validation of *in vivo* images with *ex vivo* IHC. We have shown that learning-based segmentation, via a random forest classifier, best accounted for the artefacts present in mesoscopic PAI, providing a robust segmentation of blood volume at depth in 3D and a good approximation of vessel network structure. Despite the promise of the learning-based approach to account for depth-dependent variation in SNR, auto-thresholding with vesselness filtering more accurately represents statistical and topological characteristics in the superficial blood vessels as it better preserves vessel lengths. Therefore, when quantifying PA images, users need to consider the relative importance of each descriptor as the choice of segmentation method can directly impact the resulting analyses. We have highlighted the potential of statistical and topological analyses to provide a detailed parameterisation of tumour vascular networks, from classic statistical descriptors such as vessel diameters and lengths to more complex descriptors of network topology characterising vessel connectivity and loops. Our results further underscore the potential of photoacoustic mesoscopy as a tool to provide biological insight into studying vascular network *in vivo* by providing life scientists with a readily deployable and cross-validated pipeline for data analysis.

## MATERIALS AND METHODS

### Generating ground truth vascular architectures in silico

To generate an *in silico* ground truth vascular network, we utilised Lindenmayer systems (L-Systems, see Supplementary Figure 5) (Lindenmayer, 1968). L-Systems are language-theoretic models that were originally developed to model cellular interactions but have been extended to model numerous developmental processes in biology (Rozenberg & Arto Salomaa, 1992). Here, we apply L-Systems to generate realistic, 3D vascular architectures (Galarreta-Valverde, 2012; Galarreta-Valverde et al., 2013) (referred to as L-nets) and corresponding binary image volumes. A stochastic grammar was used (Galarreta-Valverde, 2012) to create a string that was evaluated using a lexical and syntactic analyser to build a graphical representation of each L-net. To transfer the L-net to a discretised binary image volume, we used a modified Bresenham’s algorithm (Bresenham, 1965) for 3D to create a vessel skeleton. Voxels within a vessel volume were then identified using the associated vessel diameter for each centreline (Supplementary Figure 5).

### Photoacoustic image simulation of synthetic ground truths

To test the accuracy of the segmentation pipelines, the L-nets were then used to simulate *in vivo* photoacoustic vascular networks embedded in muscle tissue using the Simulation and Image Processing for Photoacoustic Imaging (SIMPA) python package (SIMPA v0.1.1, https://github.com/CAMI-DKFZ/simpa) (Janek Gröhl, Kris K. Dreher, Melanie Schellenberg, Alexander Seitel, 2021) and the k-Wave MATLAB toolbox (k-Wave v1.3, MATLAB v2020b, MathWorks, Natick, MA, USA) (Treeby & Cox, 2010). Planar illumination of the L-nets on the XY plane was achieved using Monte-Carlo eXtreme (MCX v2020, 1.8) simulation on the L-net computational grid of size 10.24 x 10.24 x 2.80 mm^3^ with 20 μm isotropic resolution. The optical forward modelling was conducted at 532 nm using the optical absorption spectrum of 50% oxygenated haemoglobin for vessels (an approximation of tumour vessel oxygenation based on previously collected photoacoustic data (Quiros-Gonzalez et al., 2018) and of water for muscle. Next, 3D acoustic forward modelling was performed on the illuminated L-nets assuming a speed of sound of 1500 ms^-1^ in k-Wave. The photoacoustic response of the illuminated L-nets was measured with a planar array of sensors positioned on the surface of the XY plane with transducer elements of bandwidth central frequency of 50 MHz (100% bandwidth) and using a 1,504 time steps, where a time step is 5×10^-8^ Hz^-1^). Finally, the 3D initial PA wave-field was reconstructed using fast Fourier transform-based reconstruction (Treeby & Cox, 2010), after adding uniform gaussian noise on the collected wave-field.

### String phantom

We used a string phantom as a ground truth structure (see **Supplementary Materials**). The agar phantom was prepared as described previously (Joseph et al., 2017) including intralipid (I141-100ML, Merck, Gillingham, UK) to mimic tissue-like scattering conditions. Red-coloured synthetic fibres (Smilco, USA) were embedded at three different depths defined by the frame of the phantom to provide imaging targets with a known diameter of 126 μm. The top string was positioned at 0.5 mm from the agar surface, the middle one at 1 mm, and the bottom one at 2 mm, as shown in Supplementary Figure 2.

### Animals

All animal procedures were conducted in accordance with project and personal licences, issued under the United Kingdom Animals (Scientific Procedures) Act, 1986 and approved locally under compliance forms CFSB1567 and CFSB1745. For *in vivo* vascular tumour models, cryopreserved breast PDX tumour fragments in freezing media composed of heat-inactivated foetal bovine serum (10500064, Gibco^TM^, Fisher Scientific, Göteborg Sweden) and 10% dimethyl sulfoxide (D2650, Merck) were defrosted at 37°C, washed with Dulbecco’s Modified Eagle Medium (41965039, Gibco) and mixed with matrigel (354262, Corning®, NY, USA) before surgical implantation. One estrogen receptor negative (ER-, n=6) PDX model and one estrogen receptor positive (ER+, n=8) PDX model were implanted subcutaneously into the flank of 6-9 week-old NOD scid gamma (NSG) mice (#005557, Jax Stock, Charles River, UK) as per standard protocols (Bruna et al., 2016). Once tumours had reached ∼1cm mean diameter, tumours were imaged and mice sacrificed afterwards, with tumours collected in formalin for IHC.

### Photoacoustic imaging

Mesoscopic PAI was performed using the raster-scan optoacoustic mesoscopy (RSOM) Explorer P50 (iThera Medical GmbH, Munich, Germany). The system uses a 532 nm laser for excitation. Two optical fibre bundles are arranged either side of a transducer, which provide an elliptical illumination beam of approximately 4 mm x 2 mm in size. The transducer and lasers collectively raster-scan across the field-of-view. A high-frequency single-element transducer with a centre frequency of 50 MHz (>90% bandwidth) detects ultrasound. The system achieves a lateral resolution of 40 μm, an axial resolution of 10 μm and a penetration depth of up to ∼3 mm (Omar et al., 2013).

For image acquisition of both phantom and mice, degassed commercial ultrasound gel (AquaSonics Parker Lab, Fairfield, NJ, USA) was applied to the surface of the imaging target for coupling to the scan interface. Images were acquired over a field of view of 12 × 12 mm^2^ (step size: 20 μm) at either 100% (phantom) or 85% (mice) laser energy and a laser pulse repetition rate of 2 kHz (phantom) or 1 kHz (mice). Image acquisition took approximately 7 min. Animals were anaesthetised using 3-5% isoflurane in 50% oxygen and 50% medical air. Mice were shaved and depilatory cream applied to remove fur that could generate image artefacts; single mice were placed into the PAI system, on a heat-pad maintained at 37°C. Respiratory rate was maintained between 70-80 bpm using isoflurane (∼1-2% concentration) throughout image acquisition.

### Segmentation and extraction of structural and topological vascular descriptors

All acquired data were subjected to pre-processing prior to segmentation, skeletonisation and structural analyses of the vascular network, with an optional step of vesselness filtering also tested (Figure 1). Prior to segmentation, data were filtered in the Fourier domain in XY plane to remove reflection lines, before being reconstructed using a backprojection algorithm in viewRSOM software (v2.3.5.2 iThera Medical GmbH) with motion correction for *in vivo* images with a voxel size of 20 x 20 x 4 μm^3^ (X,Y,Z). To reduce background noise and artefacts from the data acquisition process, reconstructed images were subjected to a high-pass filter, to remove echo noise, followed by a Wiener filter in MATLAB (v2020b, MathWorks, Natick, MA, USA) to remove stochastic noise. Then, a built-in slice-wise background correction (Sternberg, 1983) was performed in Fiji(Schindelin et al., 2012) to achieve a homogenous background intensity (see exemplars of each pre-processing step in Supplementary Figure 6).

### Image segmentation using auto-thresholding or a random forest classifier

Using two common tools adopted in the life sciences, we tested both a rule-based moment preserving thresholding method (included in Fiji v2.1.0) and a learning-based segmentation method based on random forest classifiers (with ilastik v1.3.3 (Berg et al., 2019)). These popular packages were chosen to enable widespread application of our findings. Moment preserving thresholding, referred to as *auto-thresholding* (AT) for the remainder of this work, computes the intensity moments of an image and segments the image while preserving these moments (Tsai, 1985). Training of the random forest (RF) backend was performed on 3D voxel features in labelled regions, including intensity features, as with the AT method, combined with edge filters, to account for the intensity gradient between vessels and background, and texture descriptors, to discern artefacts in the background from the brighter and more uniform vessel features, each evaluated at different scales (up to a sigma of 5.0).

A key consideration in the machine learning-based segmentation is the preparation of training and testing data (Supplementary Table 2). For the *in silico* ground truth L-net data, all voxel labels are known. All vessel labels were used for training, however, only partial background labels were supplied to minimise computational expense by labelling the 10 voxel radius surrounding all vessels as well as 3 planes parallel to the Z-axis (edges and middle) as background (Supplementary Figure 7A,B). For the phantom data, manual segmentation of the strings from background was performed to provide ground truth. Strings were segmented in all slices on which they appeared and background was segmented tightly around the string (Supplementary Figure 7C). For the *in vivo* tumour data, manual segmentation of vessels was made by a junior user (TLL) supervised by an experienced user (ELB), including images of varying signal-to-noise ratio (SNR) to increase the robustness of the algorithm for application in a range of unseen data. Up to 10 XY slices per image stack in the training dataset were segmented with pencil size 1 at different depths to account for depth-dependent SNR differences (Supplementary Figure 7D).

Between pre-processing and segmentation, feature enhancement was tested as a variable in our segmentation pipeline. In Fiji, we adapted a modified version of Sato filtering (α=0.25) (Sato et al., 1998) to calculate vesselness from Hessian matrix eigenvalues (Frangi et al., 1998) across multiple scales. Five scales in a linear Gaussian normalized scale space were used, from which the maximal response was measured to produce the final vesselness filtered images (20, 40, 60, 80, and 100 μm) (Sato et al., 1998).

Finally, all segmented images (either from Fiji or ilastik) were passed through a built-in 3D median filter in Fiji, to remove impulse noises (Supplementary Figure 8). To summarise the pipeline (Figure 1), the methods under test for all datasets were:

1. Auto-thresholding using a moment preserving method (AT);
2. Auto-thresholding using a moment preserving method with vesselness filtering pre-segmentation (AT+VF);
3. Random forest classifier (RF);
4. Random forest classifier with vesselness filtering pre-segmentation (RF+VF).

Computation times are summarised in Supplementary Table 3.

### Extracting tumour ROIs using a 3D CNN

To analyse the tumour data in isolation from the surrounding tissue required delineation of tumour regions of interest (ROIs). To achieve this, we trained a 3D convolutional neural network (CNN) to fully automate extraction of tumour ROIs from PAI volumes. The 3D CNN is based on the U-Net architecture (Ronneberger et al., 2015) extended for volumetric delineation (Çiçek et al., 2016). Details on the CNN architecture and training are provided in the **Supplementary Materials** and Supplementary Figures 9-10.

### Network Structure and Topological Data Analysis

Topological data analysis (TDA) of the vascular networks was performed using previously reported software that performs TDA and structural analyses on vasculature (Chung et al., 2019; Stolz et al., 2020). Prior to these analyses, segmented image volumes were skeletonised using the open-source package Russ-learn (Bates, 2017, 2018). Our vascular descriptors comprised a set of statistical descriptors: vessel diameters and lengths, vessel tortuosity (sum-of-angles measure, SOAM) and curvature (chord-to-length ratio, CLR), In addition, the following descriptors were used to define network topology: the number of connected components (Betti number β_0_) and looping structures (1D holes, Betti number β_1_). Full descriptions of the vascular descriptors are provided in Supplementary Table 1 while outputs are shown in Supplementary Tables 4-7.

### Immunohistochemistry

For *ex vivo* validation, formalin-fixed paraffin-embedded (FFPE) tumour tissues were sectioned. Following deparaffinising and rehydration, IHC was performed for the following antibodies: CD31 (anti-mouse 77699, Cell signalling, London, UK), α-smooth muscle actin (ASMA) (anti-mouse ab5694, abcam, Cambridge, UK), carbonic anhydrase-IX (CAIX) (anti-human AB1001, Bioscience Slovakia, Bratislava, Slovakia) at 1:100, 1:500 and 1:1000, respectively, using a BOND automated stainer with a bond polymer refine detection kit (Leica Biosystems) and 3,3’-diaminobenzadine as a substrate. Stained FFPE sections were scanned at 20x magnification using an Aperio ScanScope (Leica Biosystems, Milton Keynes, UK) and analysed using ImageScope software (Leica Biosystems) or HALO Software (v2.2.1870, Indica Labs, Albuquerque, NM, USA). ROIs were drawn over the whole viable tumour area and built-in algorithms customised to analyse the following: CD31 positive area (µm^2^) normalised to the ROI area (µm^2^) (referred to as CD31 vessel area), area of CD31 positive pixels (µm^2^) colocalised on adjacent serial section with ASMA positive pixels/CD31 positive area (µm^2^) (reported as ASMA vessel coverage (%)) and CAIX positive pixel count per total ROI pixel count (reported as CAIX (%)).

### Statistical analysis

Statistical analyses were conducted using Prism (v9, GraphPad Software, San Diego, CA, USA) and R (v4.0.1(R Core, 2021), R Foundation, Vienna, Austria). We used the mean square error and R-squared statistics to quantify the accuracy and strength of the relationship between the segmented networks to the ground truth L-nets. For each outcome of interest, we predicted the ground truth (on a scale compatible with the normality assumption according to model checks) by means of each method estimates through a linear model. As model performance statistics are typically overestimated when assessing the model fit on the same data used to estimate the model parameters, we used bootstrapping (R = 500) to correct for the optimism bias and obtain unbiased estimates (Harrell, 2016). Bland-Altman plots were produced for each paired comparison of segmented volume to the ground truth volume in L-nets and associated bias and limits of agreement (LOA) are reported. For L-nets, F1 scores were calculated (Dice, 1945). PAI quality pre-segmentation was quantified by measuring SNR, defined as the mean of signal over the standard deviation of the background signal. Comparisons of string volume, as well as SNR, were completed using one-way ANOVA with Tukey multiplicity correction.

For each outcome of interest, *in vivo* data was analysed as follows: A linear mixed effect model was fitted on a response scale (log, square root or cube root) compatible with the normality assumption according to model checks with the segmentation methods as a 4-level fixed predictor and animal as random effect, to take the within mouse dependence into account. Noting that the residual variance was sometimes different for each segmentation group, we also fitted a heteroscedastic linear mixed effect allowing the variance to be a function of the segmentation group. The results of the heteroscedastic model were preferred to results of the homoscedastic model when the likelihood ratio test comparing both models led to a p-value <0.05. Two multiplicity corrections were performed to achieve a 5% family-wise error rate for each dataset: For each outcome, a parametric multiplicity correction on the segmentation method parameters was first used (Bretz et al., 2010). A conservative Bonferroni p-value adjustment was then added to it to account for the number of outcomes in the entire *in vivo* dataset. The following pairwise comparisons were considered: AT vs. AT+VF, AT vs. RF, RF vs. RF+VF and AT+VF vs. RF+VF. Comparisons of our vascular descriptors between ER- and ER+ tumours were completed with an unpaired student’s t-test. All p-values <0.05 after multiplicity correction were considered statistically significant.

### Code Availability

Code to generate synthetic vascular trees (LNets) is available on GitHub (https://github.com/psweens/V-System). *In silico* photoacoustic simulations were performed using the SIMPA toolkit (https://github.com/CAMI-DKFZ/simpa). Both the trained 3D CNN to extract tumour ROIs from RSOM images (https://github.com/psweens/Predict-RSOM-ROI) and vascular TDA package are available on GitHub (https://github.com/psweens/Vascular-TDA).

### Data Availability

Exemplar datasets for the *in silico*, phantom, and *in vivo* data can be found at https://doi.org/10.17863/CAM.78208. The authors declare that all data supporting the findings of this study is available upon request.

## ACKNOWLEDGEMENTS

ELM, PWS, TLL, JG, LH, DLC, and SEB acknowledge the support from Cancer Research UK under grant numbers C14303/A17197, C9545/A29580, C47594/A16267, C197/A16465, C47594/A29448, and Cancer Research UK RadNet Cambridge under the grant number C17918/A28870. PWS acknowledges the support of the Wellcome Trust and University of Cambridge under the grant number RG89305. TLL is supported by the Cambridge Trust. LH is funded from NPL’s MedAccel programme financed by the Department of Business, Energy and Industrial Strategy’s Industrial Strategy Challenge Fund. BJS, HMB, and HAH are members of the Centre for Topological Data Analysis, funded by the EPSRC grant (EP/R018472/1). We thank the Cancer Research UK Cambridge Institute Biological Resources Unit, Imaging Core, Histopathological Core, Preclinical Genome Editing Core, Light Microscopy and Research Instrumentation and Cell Services for their support in conducting this research. Particular thanks go to Cara Brodie in the Histopathology Core for analyses support. ELB and SEB would like to thank Prof. Carlos Caldas, Dr Alejandra Bruna and Dr Wendy Greenwood for providing PDX tissue from their biobank at the CRUK Cambridge Institute and for assisting in the establishment of a sub-biobank that contributed the *in vivo* data presented in this manuscript.

## AUTHOR CONTRIBUTIONS

Conceptualization: PWS, ELB, TLL, SEB

Methodology: PWS, ELB, LH, TLL, ZH, SEB

Software: PWS, BJS, JG, TLL, ZH

Validation: PWS, ELB, LH, TLL

Formal Analysis: PWS, ELB, TLL

Investigation: PWS, ELB, LH, TLL

Resources: HAH, HMB

Data Curation: PWS, JG, TLL

Writing – original draft: PWS, ELB, LH, DLC, TLL, SEB

Writing – review & editing: PWS, ELB, JG, LH, BJS, HAH, HMB, TLL, SEB

Visualisation: PWS, ELB, TLL

Supervision: SEB

Project Administration: PWS, SEB

Funding Acquisition: PWS, TLL, SEB

## Supplementary information

### Supplementary Materials and Methods

#### 1. String phantom preparation

The string phantom used in this study was prepared by mixing 1.5 g agarose (Fluka Analytical, 05039-500G) in 97.3 mL deionised water in a glass media bottle and heated in a microwave until the solution turned clear. After cooling down the solution to 60°C, 2.08 mL of pre-warmed intralipid was added to generate a reduced scattering coefficient of 5.0 cm^-1^ according to a previously characterised recipe(Joseph et al., 2017). The mixture was poured into a 3D-printed phantom mould, which was designed in Autodesk Fusion 360 (San Rafael, CA, USA) and printed using an Anet A6 Printer with polylactic acid (PLA PRO 1.75mm Fluorescent Yellow PLA 3D Printer Filament, 832-0254, RS Components, UK) as a base material. Supplementary Figure 2 shows the phantom mould with and without agar.

#### 2. 3D CNN for ROI delineation

##### 2.1. Preparation of training data

Image volumes consist of a series of 8-bit grayscale *Tiffs* (no compression) of 600 x 600 pixels in the XY-plane and a stack of 700 images in Z, with anisotropic voxels of size 20 x 20 x 4 μm^3^. Our dataset has a total of 166 PAI volumes, each paired with a corresponding binary semi-manually-annotated volume, where a voxel value of 0 and 255 indicates the background or tumour ROIs, respectively. The annotated volumes were generated by an experienced user, who first identified the top and bottom image containing the tumour in Z. Within these upper and lower bounds, ROIs were manually drawn in the XY plane on approximately 4 image slices. Bound by these data, a convex hull was extrapolated to approximate the ROI in the remaining image slices.

Prior to training, image volumes and binary masks were downsampled to an isotropic volume of 256 x 256 x 256 voxels to fit into computer memory. Data was locally standardised and normalised to a pixel range between 0 and 1 and the volumes randomly partitioned into training, validation, and testing subsets. Here, ∼5% of images were allocated for testing, with the remaining portion split 80:20 for training and validation respectively (8 / 126 / 32 image volumes, respectively).

##### 2.2 Neural Network Architecture for ROI delineation

The 3D CNN is based on the U-Net architecture(Ronneberger et al., 2015) extended for volumetric delineation(Çiçek et al., 2016). The structure consists of an encoder, which extracts spatial features from a 3D image volume, and a decoder, which constructs a segmentation map from these features (Supplementary Figure 10). The network architecture consists of five convolutional layers. The encoder path contains two 3 x 3 x 3 convolutions followed by a rectified linear unit (ReLU) activation for faster convergence and accuracy(Çiçek et al., 2016). Each ReLU activation is followed by 2 x 2 x 2 max pooling with strides of two in each dimension. For the 3^rd^, 4^th^ and 5^th^ layers, dropout is applied to reduce segmentation bias and ensure segmentation is performed utilising high-level features that may not have been considered in our semi-manual ROI annotations.

The decoder path consists of two 3 x 3 x 3 deconvolutions of strides of 2 in each dimension, followed by 3 x 3 x 3 convolutions, batch normalisation and ReLU activation. High-resolution features were provided via shortcut connections from the same layer in the encoder path. The final layer applied an additional 1 x 1 x 1 convolution followed by sigmoid activation to ensure the correct number of output channels and range of pixel values [0, 1]. The input layer is designed to take *n* grayscale (one channel) tumour volumes as input with a pre-defined volume (128 x 128 x 128 voxels in X, Y, Z-direction used here). The U-Net binary mask prediction contains an equal number of voxels as the input. The CNN was implemented in Keras(Chollet & Others, 2015) with the Tensorflow framework(Abadi et al., 2015). The model was trained and tested on a Dell Precision 7920 with a Dual Intel Xeon Gold 5120 CPU with 128 GB RAM and a NVIDIA Quadro GV100 32 GB GPU.

##### 2.3. Hyperparameter Optimisation

Hyperparameters were optimised and evaluated using Talos(*Autonomio Talos*, 2019), a fully-automated hyperparameter tuner for Keras. A random search optimisation strategy was deployed using the quantum random method. Here, a probabilistic reduction scheme was used to reduce the number of parameter permutations by removing poorly performing hyperparameter configurations from the remaining search space after a predefined interval. The number of filters used ranged from 16 in the 1^st^ layer to 512 in the 5^th^. Dropout at a rate of 0.2 was applied in the 3^rd^, 4^th^ and 5^th^ layers. A Glorot uniform initialiser was used for all convolution and deconvolution layers. The model was trained using an Adam optimiser with learning and decay rates of 10^-5^ and 10^-8^, respectively, and the dice coefficient (F1)(Crum et al., 2006) used as the loss function.

##### 2.4. U-Net Training & Predictions

Training was performed with a batch size of 3 image volumes for a total of 120 epochs (Supplementary Figure 11A). The fully-trained network achieved an accuracy of 88.3% and 87.3% on the training and validation sets respectively (Supplementary Figure 11B). Following training and test, we applied the CNN to the entire set of volumes to compare predictions of ROI volume to the ground truth (Supplementary Figure 11C). Blood volumes were then calculated within the predicted ROIs using the AT method and compared against the user annotations (Supplementary Figure 11D). We found a significant correlation between user annotated and predicted data for both ROI volume (Spearman’s rank correlation: r = 0.821, p < 0.0001) and blood volume (r = 0.958, p < 0.0001), indicating our CNN achieves sufficient performance against the experienced user to be applied for extracting tumours prior to testing the segmentation pipeline.

#### 3. Signal-to-noise ratio characterisation

PAI quality pre-segmentation was quantified by measuring signal-to-noise ratio (SNR), defined as the mean of signal over the standard deviation of the background signal. For *in silico* and in phantom ground truth datasets, the mean of the signal was taken within the binary ground truth masks of the images and reported for different depths.

## Supplementary Tables

**Supplementary Table 1:** Descriptions of our statistical and topological descriptors.

| Descriptor | Description |
| --- | --- |
| Connected Components, $\beta_0$ | Number of 0-dimensional topological features, *i.e.* the number of subgraphs or clusters (vascular subnetworks). Values are normalised with respect to the total number of edges per segmented image volume. |
| Loops, $\beta_1$ | Number of 1-dimensional topological features, *i.e.* the numbers of looping structures in vascular graph. Values are normalised with respect to the total number of edges per segmented image volume. |
| Sum-of-angles measure (SOAM) | The sum of angles between tangents to the curve taken at regular intervals normalised against vessel length, *i.e.* the average change in angle per unit length. |
| Chord-to-length ratio (CLR) | The ratio between the Euclidean distance connecting the two ends of a blood vessel and the length of the blood vessel, *e.g.* a straight vessel has a CLR equal to 1. |

**Supplementary Table 2:** Training and testing dataset split for random forest-based segmentation in ilastik.

| Data | Ground truth labels | Training | Testing |
| --- | --- | --- | --- |
| *In silico* | Original binary labels of L-net branches and surrounding background | 30 L-nets | 30 L-nets (data in **Figures 2 and 3**) |
| *In vitro* | Manual labelling of all XY slices containing strings and of surrounding background | 2 string phantom scans | 5 string phantom scans (data in **Figures 4 and 5**) |
| *In vivo* | Manual labelling of 10 XY slices per image at distributed depths and of surrounding background | 20 PDX tumour scans | 14 PDX tumour scans (data in **Figures 6-9**) |

**Supplementary Table 3.** Mean computation time in seconds for each segmentation method on *in silico, in vitro*, and *in vivo* data. Note: Segmentations were performed on a dual Intel Xeon E5-2623 v4 2.60 GHz quad-core processor and 64.0 GB of RAM.

| Data | AT | AT+VF | RF | RF+VF |
| --- | --- | --- | --- | --- |
| *In silico* | 7.4 | 198.1 | 191.1 | 381.8 |
| *In vitro* | 12.9 | 219.6 | 1280.0 | 1486.7 |
| *In vivo* | 38.4 | 1215.5 | 1500.7 | 2677.8 |

**Supplementary Table 4.** Absolute number of connected components for each L-Net skeleton generated from the ground truth and each segmentation method. Network names are organised based on number of recursive L-Net iterations and index, for example, ‘LNet_i4_0’ is the zeroth network of those with 4 iterations. Note, the number of known branching points is equal to number of iterations minus 3.

| Name | Ground Truth | AT | AT+VF | RF | RF+VF |
| --- | --- | --- | --- | --- | --- |
| LNet_i4_0 | 1 | 17 | 4 | 5 | 3 |
| LNet_i4_1 | 1 | 12 | 1 | 5 | 4 |
| LNet_i4_2 | 1 | 8 | 3 | 3 | 2 |
| LNet_i4_3 | 1 | 23 | 3 | 24 | 111 |
| LNet_i4_4 | 1 | 15 | 1 | 2 | 1 |
| LNet_i6_0 | 1 | 12 | 3 | 10 | 16 |
| LNet_i6_1 | 1 | 21 | 2 | 4 | 6 |
| LNet_i6_2 | 1 | 13 | 3 | 4 | 8 |
| LNet_i6_3 | 1 | 1 | 2 | 3 | 2 |
| LNet_i6_4 | 1 | 21 | 9 | 8 | 4 |
| LNet_i8_0 | 1 | 20 | 9 | 16 | 16 |
| LNet_i8_1 | 1 | 26 | 5 | 16 | 9 |
| LNet_i8_2 | 1 | 12 | 9 | 12 | 5 |
| LNet_i8_3 | 1 | 12 | 7 | 13 | 6 |
| LNet_i8_4 | 1 | 7 | 2 | 14 | 11 |
| LNet_i10_0 | 1 | 30 | 14 | 29 | 21 |
| LNet_i10_1 | 1 | 30 | 12 | 24 | 19 |
| LNet_i10_2 | 1 | 18 | 9 | 24 | 14 |
| LNet_i10_3 | 1 | 40 | 20 | 33 | 34 |
| LNet_i10_4 | 1 | 21 | 21 | 28 | 27 |
| LNet_i12_0 | 1 | 76 | 16 | 49 | 43 |
| LNet_i12_1 | 1 | 68 | 23 | 52 | 52 |
| LNet_i12_2 | 1 | 58 | 24 | 40 | 37 |
| LNet_i12_3 | 1 | 49 | 19 | 83 | 72 |
| LNet_i12_4 | 1 | 58 | 26 | 68 | 53 |
| LNet_i14_0 | 1 | 88 | 39 | 155 | 103 |
| LNet_i14_1 | 1 | 81 | 45 | 112 | 100 |
| LNet_i14_2 | 1 | 69 | 36 | 93 | 79 |
| LNet_i14_3 | 1 | 91 | 46 | 87 | 89 |
| LNet_i14_4 | 1 | 104 | 34 | 116 | 74 |

**Supplementary Table 5.** Absolute number of loops for each L-Net skeleton generated from the ground truth and each segmentation method. Network names are organised based on number of recursive L-Net iterations and index, for example, ‘LNet_i4_0’ is the zeroth network of those with 4 iterations. Note, the number of known branching points is equal to number of iterations minus 3.

| Name | Ground Truth | AT | AT+VF | RF | RF+VF |
| --- | --- | --- | --- | --- | --- |
| LNet_i4_0 | 0 | 46 | 0 | 2 | 2 |
| LNet_i4_1 | 0 | 27 | 0 | 4 | 16 |
| LNet_i4_2 | 0 | 42 | 0 | 0 | 8 |
| LNet_i4_3 | 1 | 45 | 13 | 67 | 86 |
| LNet_i4_4 | 0 | 41 | 0 | 1 | 11 |
| LNet_i6_0 | 0 | 54 | 1 | 12 | 27 |
| LNet_i6_1 | 1 | 28 | 0 | 13 | 11 |
| LNet_i6_2 | 1 | 63 | 7 | 22 | 72 |
| LNet_i6_3 | 0 | 6 | 0 | 0 | 0 |
| LNet_i6_4 | 2 | 47 | 0 | 9 | 8 |
| LNet_i8_0 | 4 | 68 | 1 | 33 | 22 |
| LNet_i8_1 | 2 | 21 | 0 | 4 | 1 |
| LNet_i8_2 | 2 | 53 | 0 | 2 | 11 |
| LNet_i8_3 | 1 | 86 | 0 | 8 | 13 |
| LNet_i8_4 | 1 | 40 | 0 | 1 | 0 |
| LNet_i10_0 | 4 | 0 | 0 | 0 | 0 |
| LNet_i10_1 | 20 | 14 | 0 | 0 | 1 |
| LNet_i10_2 | 9 | 20 | 0 | 0 | 0 |
| LNet_i10_3 | 9 | 33 | 0 | 14 | 12 |
| LNet_i10_4 | 11 | 7 | 0 | 1 | 1 |
| LNet_i12_0 | 73 | 9 | 4 | 24 | 12 |
| LNet_i12_1 | 123 | 37 | 5 | 25 | 25 |
| LNet_i12_2 | 106 | 32 | 7 | 35 | 36 |
| LNet_i12_3 | 30 | 4 | 1 | 1 | 1 |
| LNet_i12_4 | 62 | 2 | 1 | 8 | 13 |
| LNet_i14_0 | 353 | 16 | 9 | 18 | 13 |
| LNet_i14_1 | 426 | 43 | 15 | 58 | 43 |
| LNet_i14_2 | 395 | 19 | 9 | 19 | 20 |
| LNet_i14_3 | 376 | 74 | 15 | 58 | 51 |
| LNet_i14_4 | 304 | 29 | 12 | 20 | 29 |

**Supplementary Table 6.** The number of edges for each L-Net skeleton generated from the ground truth and each segmentation method. Network names are organised based on number of recursive L-Net iterations and index, for example, ‘LNet_i4_0’ is the zeroth network of those with 4 iterations. Note, the number of known branching points is equal to number of iterations minus 3.

| Name | Ground Truth | AT | AT+VF | RF | RF+VF |
| --- | --- | --- | --- | --- | --- |
| LNet_i4_0 | 3 | 160 | 4 | 19 | 10 |
| LNet_i4_1 | 3 | 122 | 3 | 24 | 59 |
| LNet_i4_2 | 3 | 144 | 3 | 5 | 32 |
| LNet_i4_3 | 7 | 177 | 49 | 247 | 420 |
| LNet_i4_4 | 3 | 136 | 3 | 9 | 38 |
| LNet_i6_0 | 15 | 217 | 18 | 63 | 122 |
| LNet_i6_1 | 18 | 134 | 12 | 55 | 60 |
| LNet_i6_2 | 19 | 222 | 34 | 94 | 236 |
| LNet_i6_3 | 15 | 37 | 14 | 15 | 14 |
| LNet_i6_4 | 21 | 167 | 11 | 50 | 43 |
| LNet_i8_0 | 65 | 267 | 34 | 172 | 130 |
| LNet_i8_1 | 65 | 139 | 19 | 77 | 57 |
| LNet_i8_2 | 69 | 227 | 35 | 68 | 85 |
| LNet_i8_3 | 62 | 315 | 33 | 85 | 92 |
| LNet_i8_4 | 62 | 148 | 18 | 44 | 37 |
| LNet_i10_0 | 238 | 114 | 46 | 113 | 101 |
| LNet_i10_1 | 251 | 161 | 54 | 116 | 111 |
| LNet_i10_2 | 196 | 164 | 43 | 100 | 86 |
| LNet_i10_3 | 179 | 241 | 64 | 197 | 183 |
| LNet_i10_4 | 253 | 138 | 69 | 151 | 138 |
| LNet_i12_0 | 687 | 211 | 97 | 332 | 282 |
| LNet_i12_1 | 669 | 283 | 103 | 296 | 296 |
| LNet_i12_2 | 698 | 310 | 130 | 388 | 349 |
| LNet_i12_3 | 794 | 188 | 84 | 182 | 197 |
| LNet_i12_4 | 662 | 150 | 79 | 249 | 246 |
| LNet_i14_0 | 2359 | 317 | 175 | 524 | 396 |
| LNet_i14_1 | 2226 | 463 | 262 | 642 | 591 |
| LNet_i14_2 | 2037 | 261 | 158 | 461 | 427 |
| LNet_i14_3 | 1707 | 459 | 174 | 506 | 461 |
| LNet_i14_4 | 1936 | 345 | 153 | 462 | 460 |

**Supplementary Table 7.** The number of nodes for each L-Net skeleton generated from the ground truth and each segmentation method. Network names are organised based on number of recursive L-Net iterations and index, for example, ‘LNet_i4_0’ is the zeroth network of those with 4 iterations. Note, the number of known branching points is equal to number of iterations minus 3.

| Name | Ground Truth | AT | AT+VF | RF | RF+VF |
| --- | --- | --- | --- | --- | --- |
| LNet_i4_0 | 4 | 131 | 8 | 22 | 11 |
| LNet_i4_1 | 4 | 107 | 4 | 25 | 47 |
| LNet_i4_2 | 4 | 110 | 6 | 8 | 26 |
| LNet_i4_3 | 7 | 155 | 39 | 204 | 445 |
| LNet_i4_4 | 4 | 110 | 4 | 10 | 28 |
| LNet_i6_0 | 16 | 175 | 20 | 61 | 111 |
| LNet_i6_1 | 18 | 127 | 14 | 46 | 55 |
| LNet_i6_2 | 19 | 172 | 30 | 76 | 172 |
| LNet_i6_3 | 16 | 32 | 16 | 18 | 16 |
| LNet_i6_4 | 20 | 141 | 20 | 49 | 39 |
| LNet_i8_0 | 62 | 219 | 42 | 155 | 124 |
| LNet_i8_1 | 64 | 144 | 24 | 89 | 65 |
| LNet_i8_2 | 68 | 186 | 44 | 78 | 79 |
| LNet_i8_3 | 62 | 241 | 40 | 90 | 85 |
| LNet_i8_4 | 62 | 115 | 20 | 57 | 48 |
| LNet_i10_0 | 235 | 144 | 60 | 142 | 122 |
| LNet_i10_1 | 232 | 177 | 66 | 140 | 129 |
| LNet_i10_2 | 188 | 162 | 52 | 124 | 100 |
| LNet_i10_3 | 171 | 248 | 84 | 216 | 205 |
| LNet_i10_4 | 243 | 152 | 90 | 178 | 164 |
| LNet_i12_0 | 615 | 278 | 109 | 357 | 313 |
| LNet_i12_1 | 547 | 314 | 121 | 323 | 323 |
| LNet_i12_2 | 593 | 336 | 147 | 393 | 350 |
| LNet_i12_3 | 765 | 233 | 102 | 264 | 268 |
| LNet_i12_4 | 601 | 206 | 104 | 309 | 286 |
| LNet_i14_0 | 2007 | 389 | 205 | 661 | 486 |
| LNet_i14_1 | 1801 | 501 | 292 | 696 | 648 |
| LNet_i14_2 | 1643 | 311 | 185 | 535 | 486 |
| LNet_i14_3 | 1332 | 476 | 205 | 535 | 499 |
| LNet_i14_4 | 1633 | 420 | 175 | 558 | 505 |

## Supplementary Figures

**Supplementary Figure 1.**
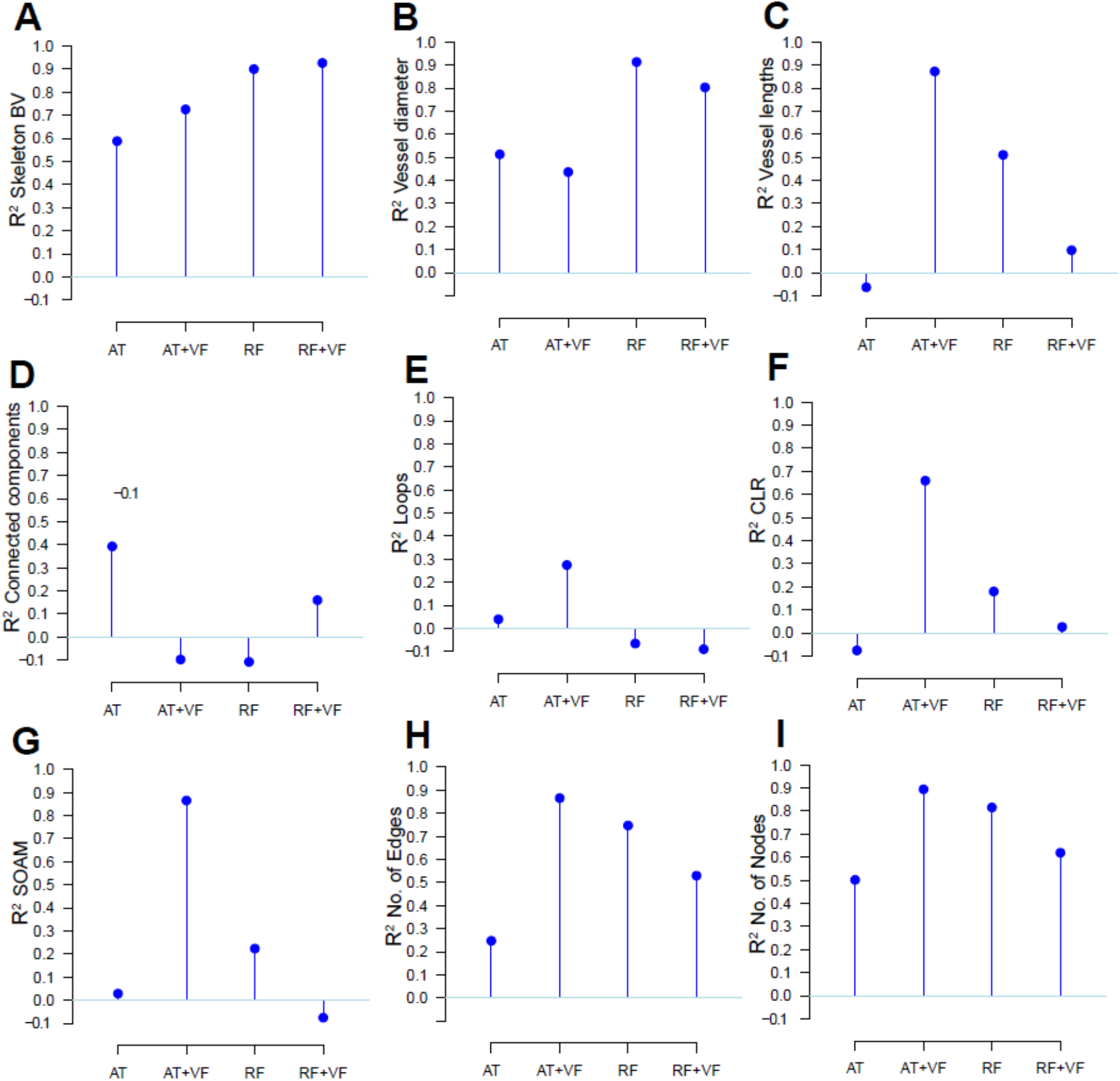
Random forest classifier segments PAI networks with high accuracy while autothresholding with vesselness filtering preserves network structure. Bar plot for R^2^ values calculated to compare the strength of relationship between the segmented networks (AT, AT+VF, RF or RF+VF) and ground-truth L-nets for the following metrics: (A) Normalised skeleton blood volume (BV), (B) Vessel diameters, µm, (C) Vessel lengths, µm, (D) Connected components, (E) Loops, (F) chord-to-length ratio (CLR), (G) sum-of-angle measure (SOAM), (H) Number of Edges and (I) Number of Nodes.

**Supplementary Figure 2.**
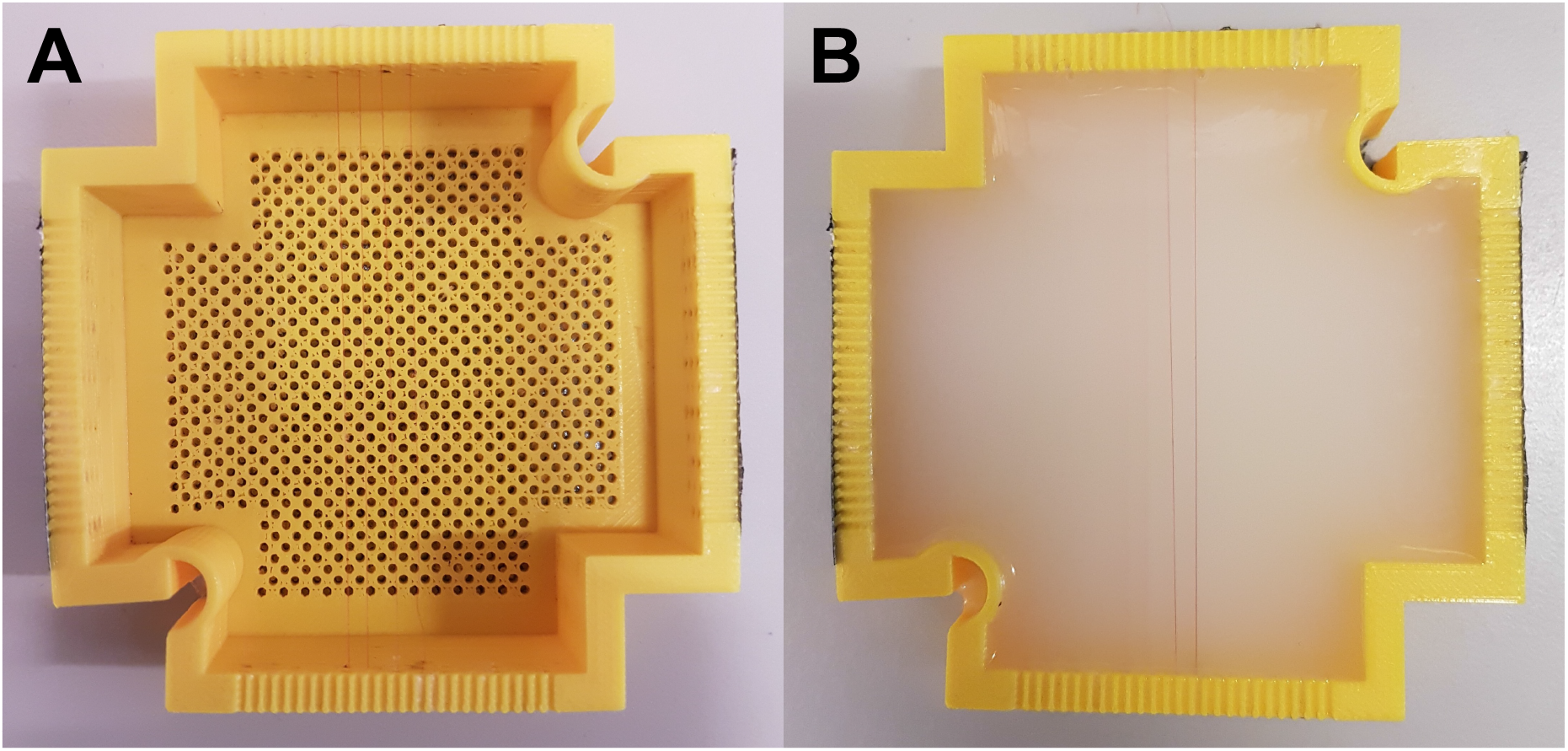
Photographs of the string phantom. (A) 3D-printed mould (7.4 x 7.4 cm, wall thickness: 4 mm) with the embedded strings and (B) with the agar gel. The top string was positioned at 0.5 mm from the agar surface, the middle one at 1 mm, and the bottom one at 2 mm depth.

**Supplementary Figure 3.**
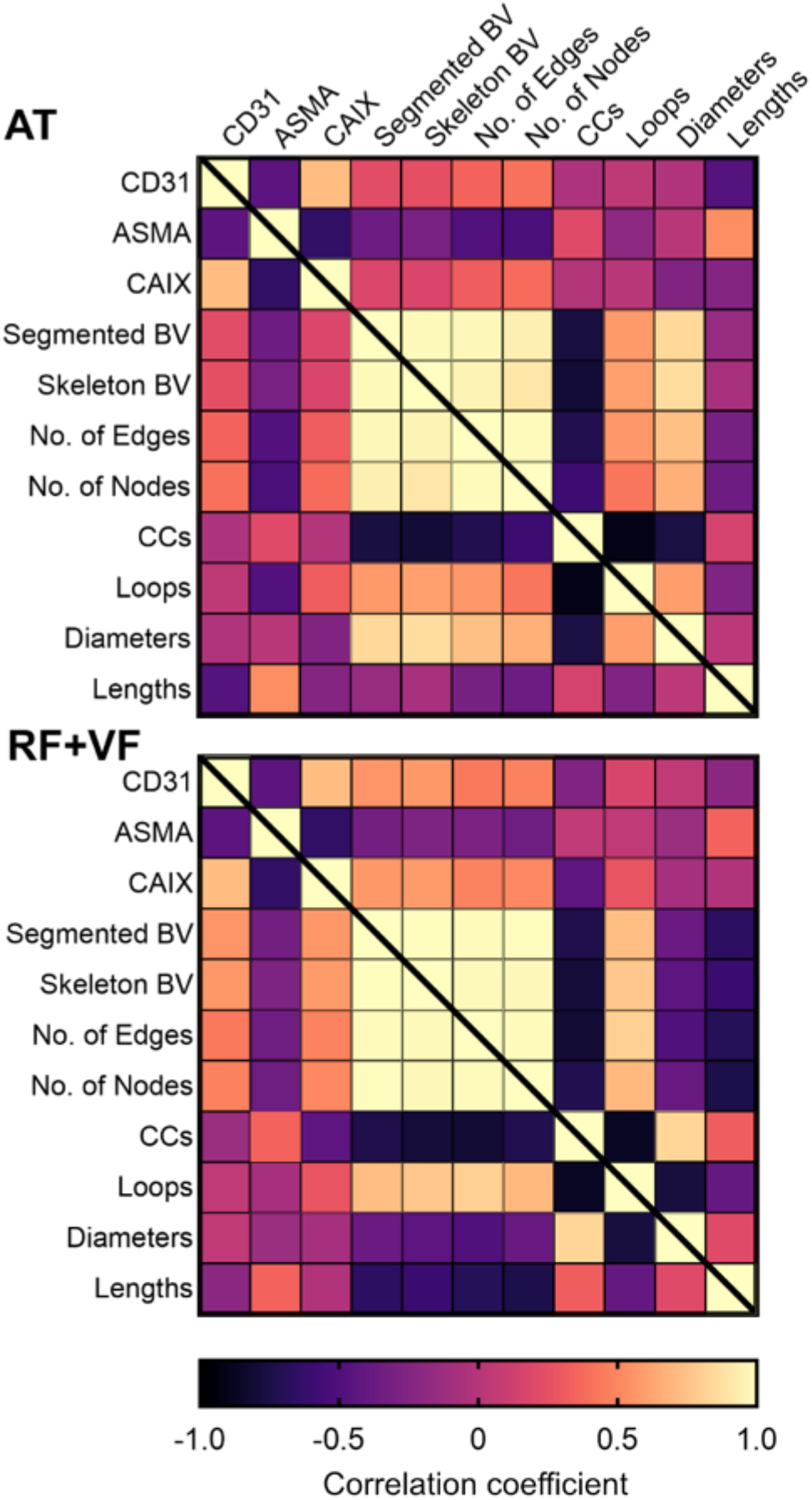
Correlation between blood volume and statistical and topological *in vivo* metrics with *ex vivo* IHC in AT and RF+VF segmented networks. Matrix of correlation coefficients for AT (top) and RF+VF (bottom) segmented networks. Pearson or spearman coefficients are used as appropriate, depending on data distribution. Note that none of the coefficients are significant for AT networks (p>0.05). For RF+VF, CD31 staining area and CAIX significantly correlated with segmented (p=0.04 and p=0.03 respectively) and skeletonised blood volume (p=0.03 for both).

**Supplementary Figure 4.**
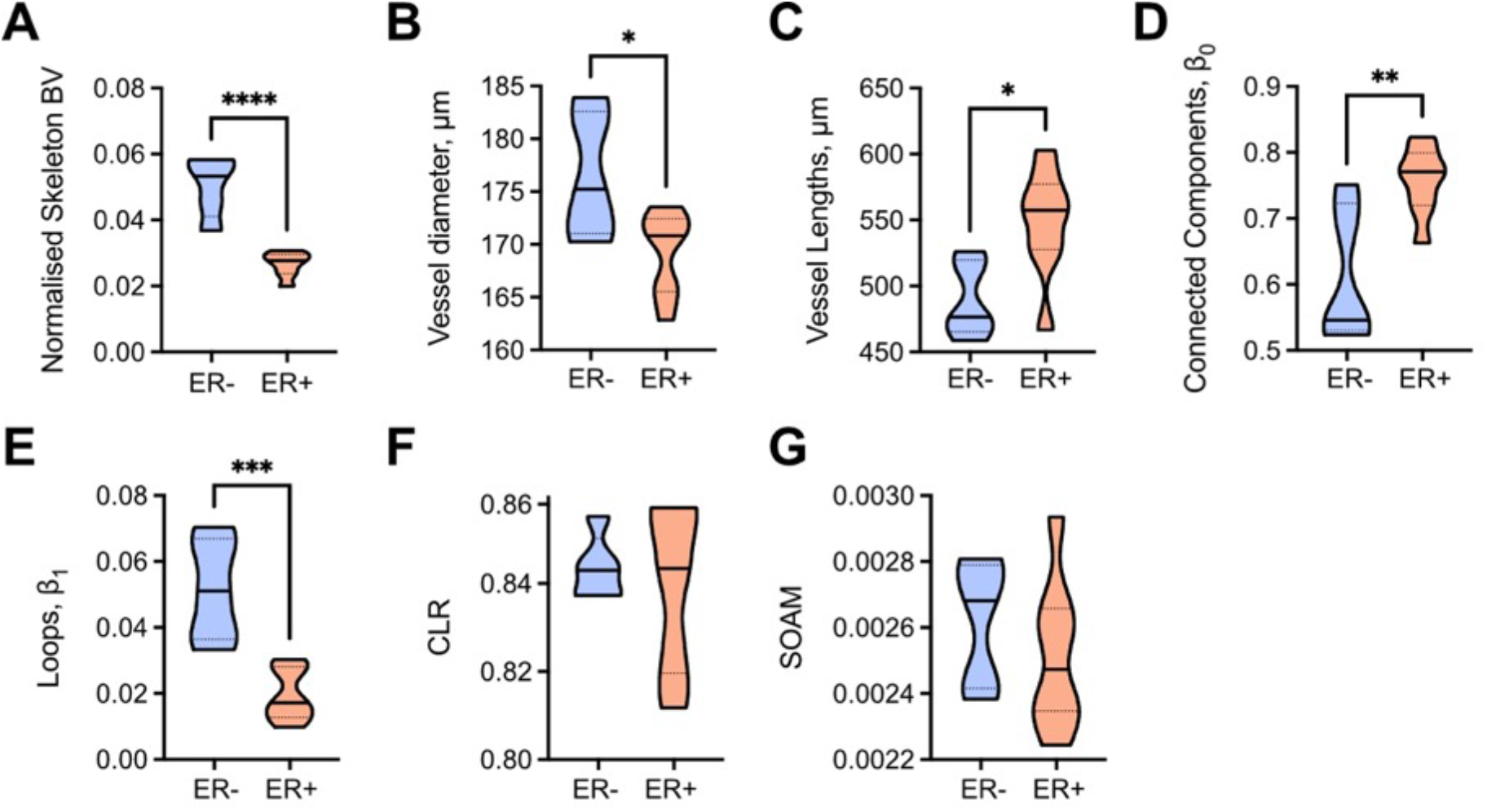
Statistical and topological analyses of AT+VF segmentation masks comparing ER- and ER+ tumours. (A-G) Abbreviations defined: blood volume (BV), chord-to-length ratio (CLR), sum-of-angle measure (SOAM). Data are represented by truncated violin plots with interquartile range (dotted black) and median (solid black). Comparisons between ER- and ER+ tumours made with unpaired t-test. *= p<0.05, **=p<0.01, ***=p<0.001, ****=p<0.0001.

**Supplementary Figure 5.**
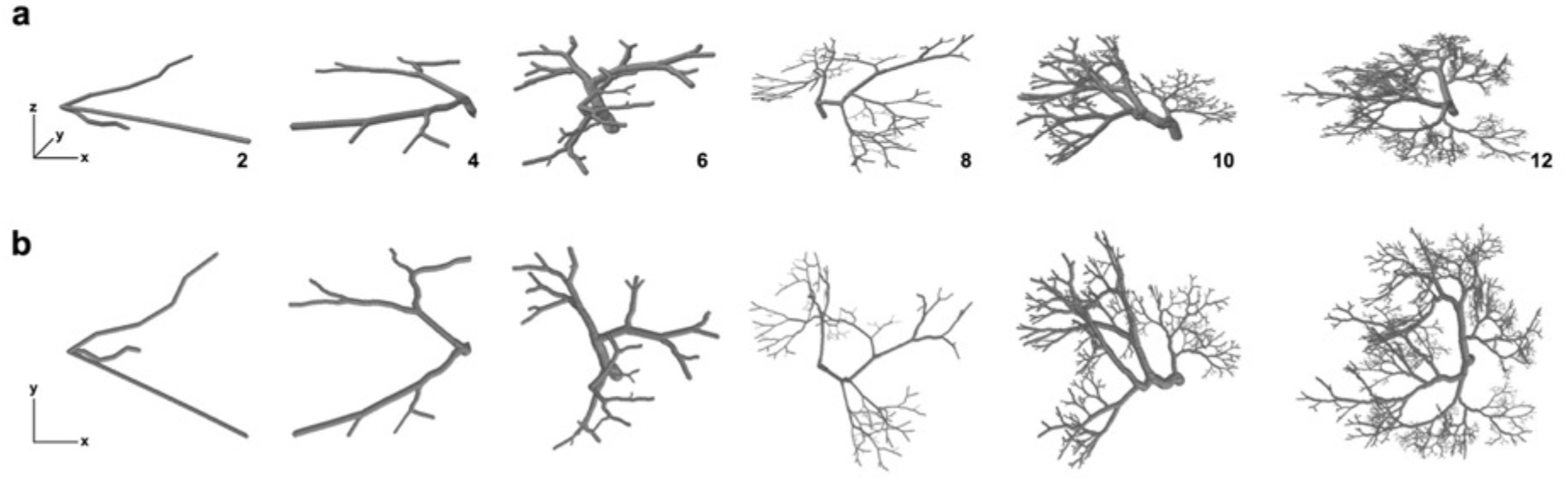
Generation of Lindenmayer System (L-System) vascular networks. (A) Segmented views of L-System vasculatures for an increasing number of branching generations (left to right; number of generations indicated). (B) Projected view in the (X,Y) plane of the architectures shown in (A).

**Supplementary Figure 6.**
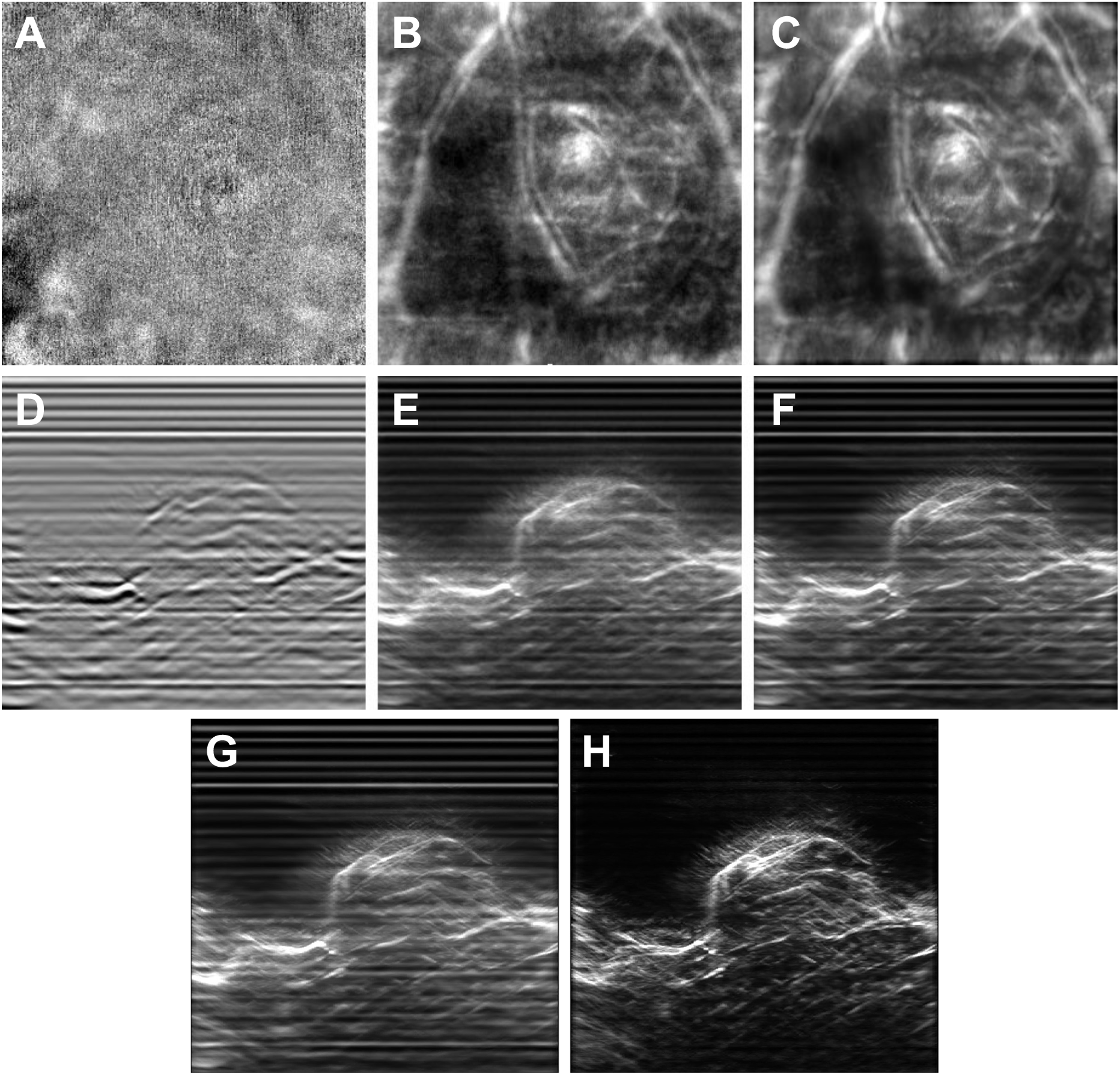
RSOM data pre-processing in MATLAB. Mean Intensity Projection 2D view of an example RSOM tumour dataset along Z axis (A-C) and Y axis (D-F) axis. From left to right: raw data (A,D), high-pass filtered data (B,E), Wiener filtered data (C,F). The images are processed sequentially through this pipeline, using high-pass filtering to remove echo noises and low-pass adaptive Wiener filtering to further remove stochastic noise in the datasets. (G) Image after MATLAB pre-processing. (H) Image after background correction with rolling ball subtraction in Fiji. The periodical horizontal line artefacts are mostly removed after background correction. All images are 6 x 6 mm.

**Supplementary Figure 7.**
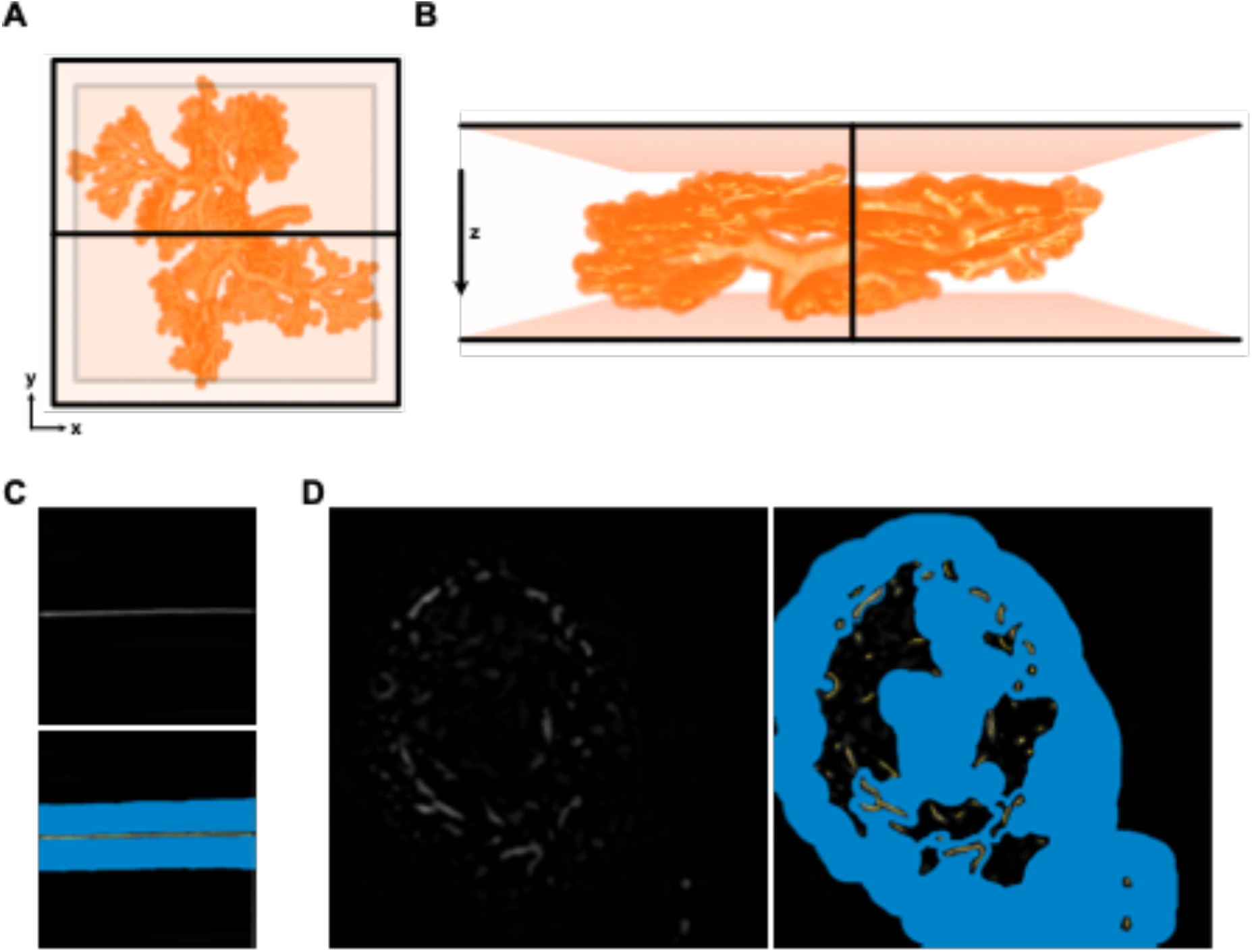
Labelling of photoacoustic data for random forest classifier training with ilastik. (A,B) Labels for the full vascular architecture of a given L-net were used for training of ilastik. The region of the L-net within 10 voxels of the vessels was labelled as background (dark orange) in addition to a three voxel thick planes (shown in black). The first was located parallel to the z-axis, with the remaining two perpendicular at the top and bottom of the image volume. (C) Labelling of string volumes and (D) of PDX tumour vessels for ilastik training. For (C) and (D) background was labelled as blue and vessels labelled as yellow on 2D slices throughout the 3D volume stack. All images are 6 x 6 mm.

**Supplementary Figure 8.**
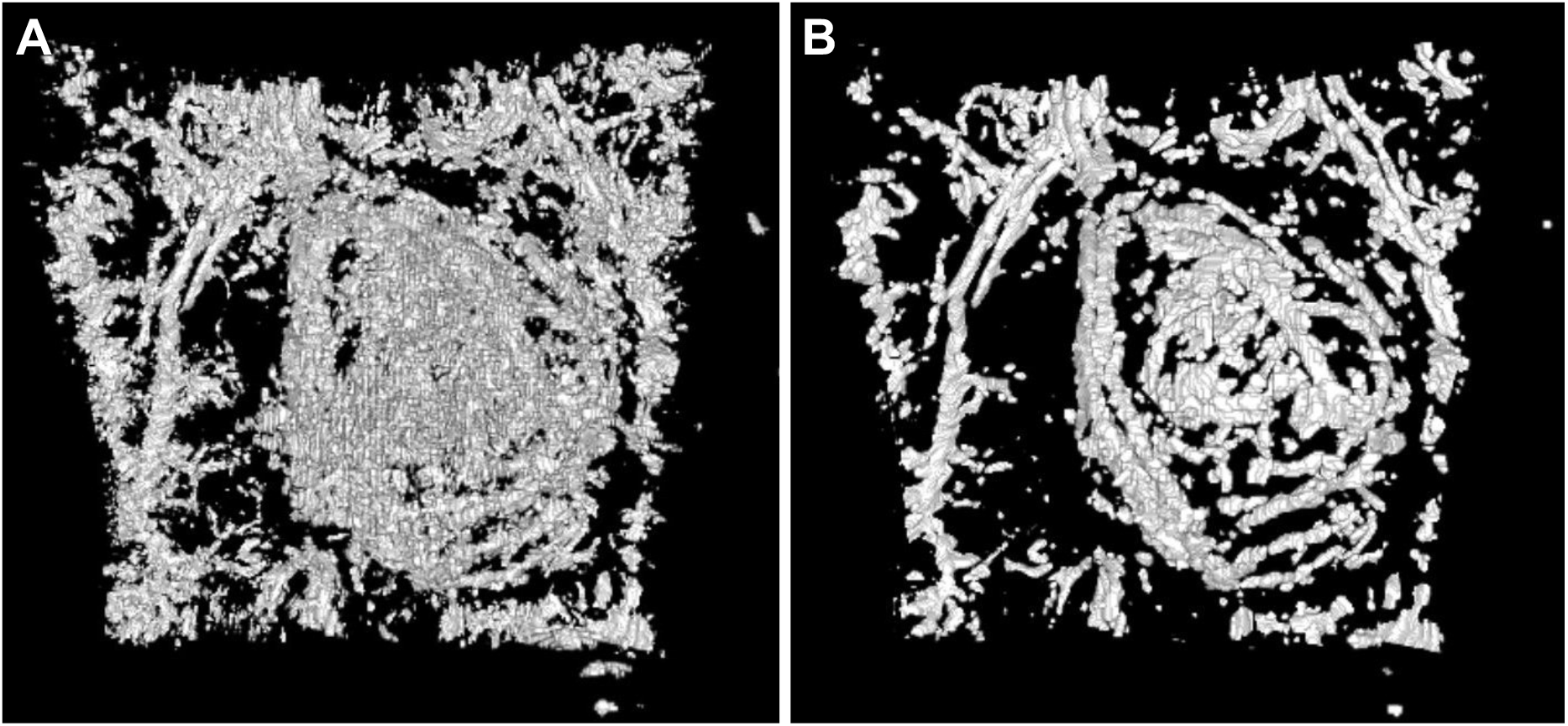
Median filtering of segmented RSOM images. A 3D rendering of the exemplar RSOM dataset (6 x 6 x 2.5 mm in X, Y and Z dimensions) used in Supplementary Figure 6 is shown. (A) Autothresholded dataset. (B) Autothresholded dataset after 3D Median filtering, to remove impulse noise.

**Supplementary Figure 9.**
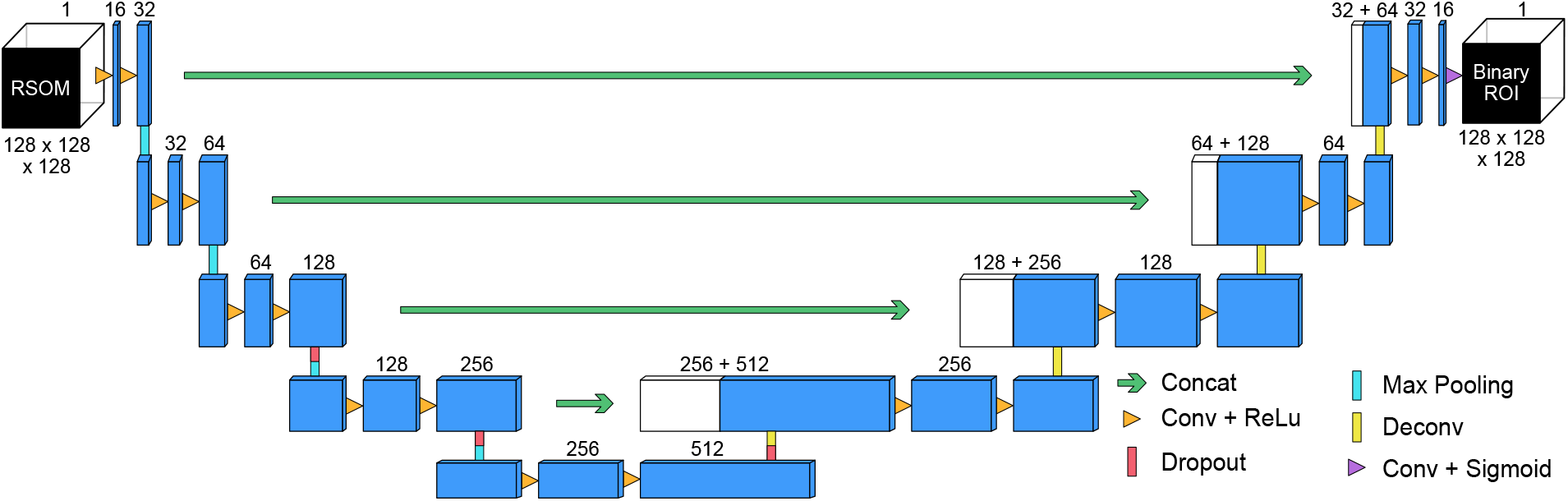
3D U-Net architecture. The blue boxes indicate feature maps with the number of channels denoted above. The input and output image volumes consist of 128 x 128 x 128 voxels. Concat = concatenation, Conv = convolution, ReLu = rectified linear unit, Deconv = deconvolution.

**Supplementary Figure 10.**
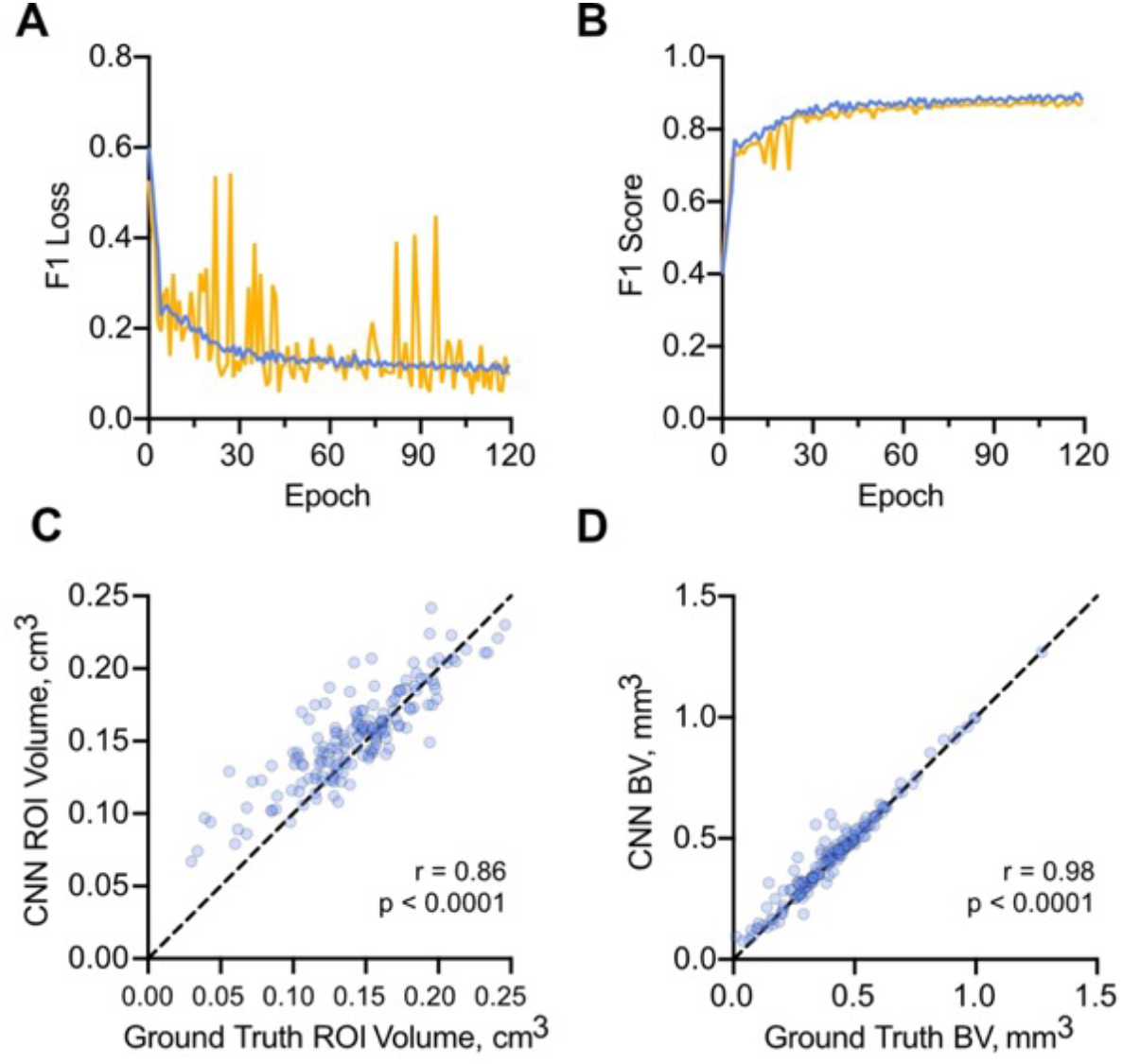
U-Net training metrics and predictions from the fully-trained architecture. Training metrics: (A) F1 loss and (B) F1 score for the training (blue) and validation (orange) datasets. (C) Region-of-interest volumes calculated from the ground truth (GT) versus the U-Net mask. (D) Computed blood volumes using the ground truth and U-Net ROI estimations from (C). Note, the lines in (C) and (D) indicate a 1-to-1 relationship, and blood volumes in (B) were calculated using our auto-thresholding segmentation method.

