## Supplementary figures and images for "Quantification of vascular networks in photoacoustic mesoscopy"

### Supplementary Movie 1

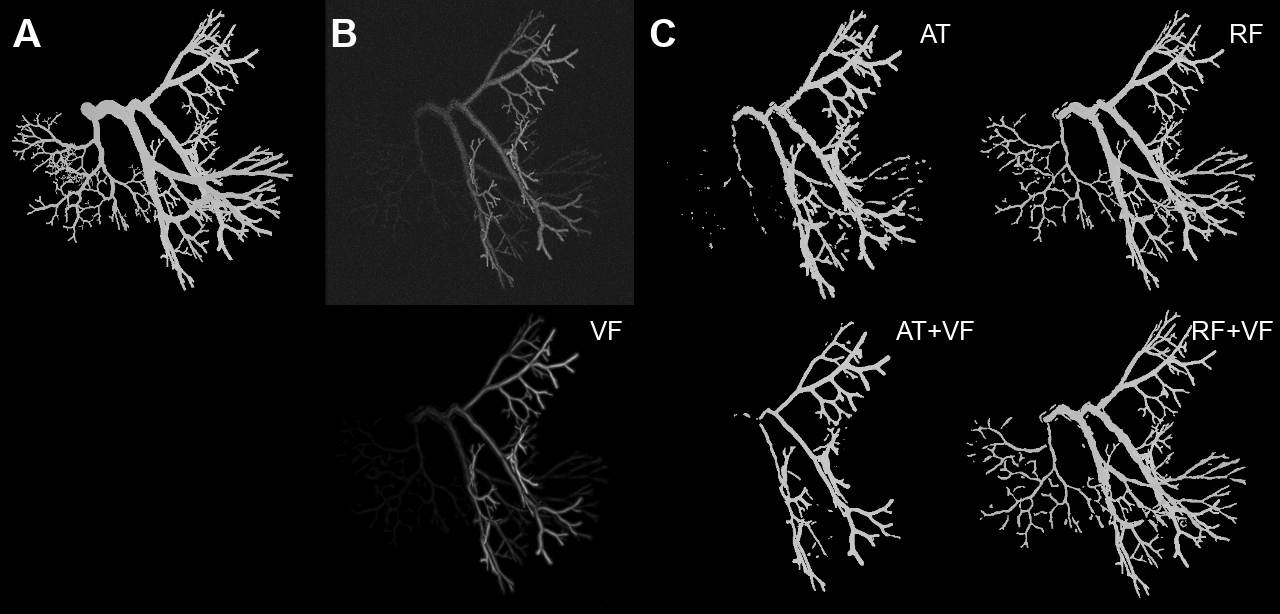

### Supplementary Movie 2

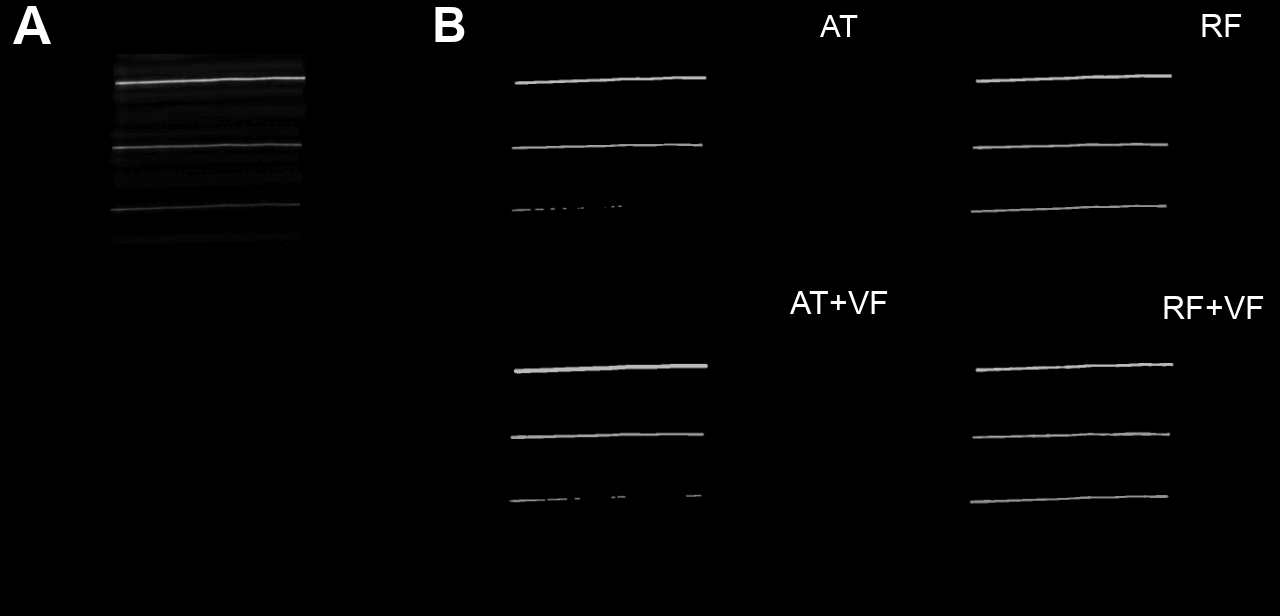

### Supplementary Movie 3

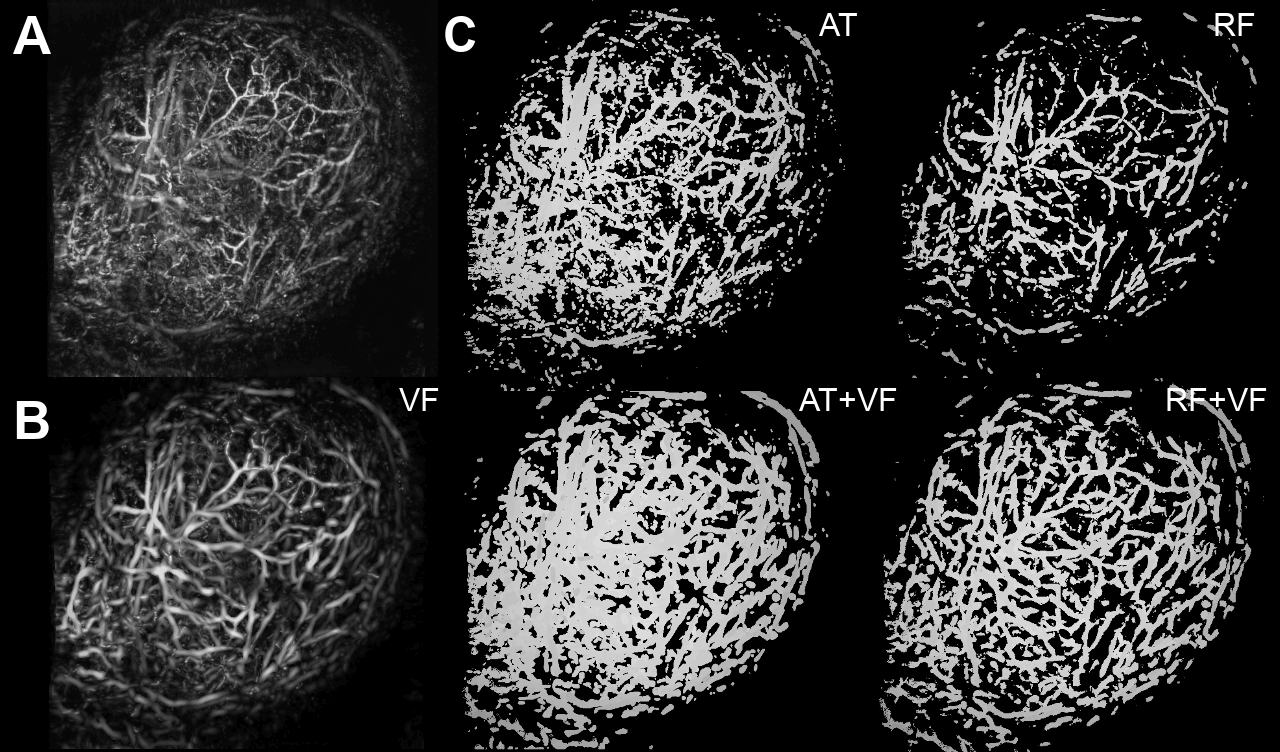
